# Characterization of human Metaxin proteins reveals functional diversification of SAM37 homologs MTX1 and MTX3

**DOI:** 10.64898/2026.03.15.711473

**Authors:** Sarah E. J. Morf, Matthew P. Challis, Sanjeev Uthishtran, Caitlin L. Rowe, Alice J. Sharpe, Natasha Kapoor-Kaushik, Senthil Arumugam, Luke E. Formosa, Kate McArthur, Michael T. Ryan

**Affiliations:** Department of Biochemistry and Molecular Biology, Biomedicine Discovery Institute, Monash University, Melbourne, VIC, Australia; Department of Anatomy and Developmental Biology, Biomedicine Discovery Institute, Monash University, Melbourne, VIC, Australia; EMBL Australia, Monash University, Melbourne, VIC, Australia; Monash Ramaciotti Centre for Cryo-Electron Microscopy, Monash University, Melbourne, VIC, Australia

**Keywords:** Mitochondria, Metaxin, MTX1, MTX2, MTX3, biogenesis, protein import, sorting and assembly machinery, SAM50

## Abstract

The biogenesis of outer mitochondrial membrane β-barrel proteins relies on the mitochondrial Sorting and Assembly Machinery (SAM) complex. In humans, the SAM complex contains SAM50 along with Metaxin (MTX) accessory subunits. MTX1 and MTX3 are homologous yet their functional similarities and differences have scarcely been investigated. Homozygous null mutations in the *MTX2* gene are linked to a rare progeroid syndrome that causes severe depletion of MTX1. Here, we uncover unique phenotypes associated with the loss of MTX1 or MTX3 in human cells. Loss of MTX1 confers a deficiency in mitochondrial volume and causes network-wide mitochondrial morphology abnormalities. MTX3 loss resulted in negligible consequences for the biogenesis of β-barrel proteins but resulted in increased mitochondrial mass. We also find that both MTX1 and MTX3 stability are dependent on the presence of MTX2, with MTX1 deficiency causing defective import and assembly. Collectively, our findings support the notion that MTX1 and MTX3 are functionally diverse homologs and are unlikely to be functionally redundant.

## Introduction

Mitochondria are host to a myriad of metabolic processes which supply the cell with energy-rich molecules and essential metabolites[1,2]. While mitochondria harbour their own genome (mitochondrial DNA; mtDNA), the majority (∼99%) of mitochondrial proteins are encoded by the nuclear genome and are translated on cytosolic ribosomes before their subsequent import into the organelle[3,4]. Distinct protein import machineries local to both the Outer Mitochondrial Membrane (OMM) and the Inner Mitochondrial Membrane (IMM) ensure correct sub-compartmental targeting of mitochondrial pre-proteins and, in many cases, their folding and maturation[5]. The OMM houses two major protein import complexes – the Translocase of the Outer Membrane (TOM) complex and the Sorting and Assembly Machinery (SAM) complex. Across species, the TOM complex facilitates the passage of nuclear-encoded mitochondrial pre-proteins from the cytosol across the OMM and into mitochondria[6]. Meanwhile, the SAM complex orchestrates the biogenesis and integration of β-barrel proteins into the OMM[7,8]. Notably, the core subunits of the SAM and TOM complexes, SAM50 and TOM40 respectively, are themselves β-barrel proteins, attesting to the bacterial ancestry of mitochondria[9]. Indeed, β-barrel proteins represent a substantial portion of the OMM proteome, highlighting the importance of the SAM complex for the maintenance of mitochondrial protein content[10].

Much of our current understanding of β-barrel protein biogenesis emanates from rigorous characterization of the β-barrel protein import pathway in yeast[9]. The yeast SAM complex is composed of three subunits, including the core subunit, SAM50, which associates with two cytosol-facing accessory subunits, SAM35 and SAM37[11]. SAM35 is a peripheral membrane protein which caps the cytosolic face of the SAM50 β-barrel and is stabilized by interactions between the SAM35 N-terminal region and cytosolic loops of the SAM50 β-barrel[11,12]. Meanwhile, SAM37 is stabilized via interactions with SAM35 on the cytosolic face of the complex and the SAM50 IMS-facing domain[11]. In humans, Metaxin 2 (MTX2) shares homology with SAM35[13]. Patients bearing homozygous null mutations in the *MTX2* gene present with a progeria-like syndrome known as Mandibuloacral dysplasia associated to *MTX2* (MADaM)[14,15]. Functional characterisation of MADaM patient fibroblasts revealed that MTX2 deficiency was associated with abnormal mitochondrial morphology, altered OXPHOS capacity, desensitization to extrinsic apoptotic stimuli and elevated baseline mitophagy[14]. Meanwhile, SAM37 has two vertebrate homologs, Metaxin 1 (MTX1) and Metaxin 3 (MTX3)[13]. The Metaxin proteins are also widely regarded as the cytosol-facing components of the Mitochondrial Intermembrane Space Bridging (MIB) complex, a ∼2.2MDa complex which tethers the OMM to the IMM via interactions between the IMS-face of SAM50 and subunits of the Mitochondrial Contact Site and Cristae Organising System (MICOS)[16,17]. Metaxins share low sequence identity with their yeast ancestors[18], and their requirement for the folding and maturation of β-barrel proteins in higher-order eukaryotes is poorly understood. Moreover, the common and diverse functions of duplicate genes, MTX1 and MTX3, are yet to be elucidated.

Here, we demonstrate that MTX1 and MTX3 have adopted discrete roles in selected mitochondrial processes and provide foundational evidence for their functional diversification. By examining the phenotypic signature associated with the loss of MTX1 or MTX3 in human cells, we uncovered both exclusive and opposing phenotypes regarding mitochondrial protein content, cellular mitochondrial volume and mitochondrial morphology arising from genetic ablation of MTX1 and MTX3. Additionally, we established a hierarchy of stability between MTX1, MTX2 and MTX3, and determined the requirement of MTX2 for the stability of MTX1 and MTX3 to be unidirectional. We examined the consequences of MTX1, MTX2 and MTX3 loss on the import and assembly kinetics of human SAM complex substrates and found clear discrepancies in the requirement of MTX1 and MTX3 for β-barrel biogenesis and maintenance of steady-state protein levels. Building upon previous knowledge defining the complexome profiles of MTX1 and MTX3[13], we affirmed the differences in the assembly states occupied by MTX1 and MTX3 and, by extension, identified unique interacting partners of each protein.

## Results

### Genetic ablation of MTX1 or MTX3 differentially affects mitochondrial protein content

To investigate the impact of MTX1, MTX2 or MTX3 loss on mitochondrial health, we used CRISPR/Cas9-mediated genome editing to establish distinct Metaxin knockout (KO) models in human osteosarcoma (U2OS) cells. The absence of MTX1 or MTX2 protein was confirmed by SDS-PAGE and immunoblotting using mitochondria isolated from MTX1^KO^ and MTX2^KO^ cells **(Fig. 1A-B)**. Due to the unavailability of suitable antibodies, MTX3^KO^ cells were validated by genotyping **(Fig. S1)**. As reported previously[14,17,19], MTX1 levels were depleted upon genetic ablation of MTX2 but were restored upon complementation with a stably-expressed N-terminal FLAG-tagged version of MTX2 (^FLAG^MTX2) **(Fig. 1B)**. Conversely, the effect of MTX1 loss on MTX2 levels was negligible **(Fig. 1A)**, and neither the levels of MTX1 or MTX2 were altered in the absence of MTX3 **(Fig. 1C)**. Across all three genotypes, the levels of SAM50, previously reported to be required for MTX1 and MTX2 stability[19], were unaltered, suggesting that the dependency of Metaxins on SAM50 for their stability is unidirectional **(Fig. 1A-C)**. Contrary to a previous report[19], we observed a decrease in TOM40 abundance in the absence of MTX1 and MTX2 which was corrected upon re-expression of ^FLAG^MTX1 or ^FLAG^MTX2 respectively **(Fig. 1A-B)**. Since the loss of TOM40 was independent of any changes to SAM50, we reasoned that MTX1 and MTX2, but not MTX3, may function as accessories to the SAM complex to maintain TOM40 levels. To investigate the requirement of MTX1, MTX2 or MTX3 for the stability of the MICOS and SAM complexes, we performed Blue Native (BN)-PAGE analysis using mitochondria isolated from MTX1^KO^, MTX2^KO^ and MTX3^KO^ cells **(Fig. 1D-F)**. We noted that the SAM complex resolved as a species of ∼200kDa, consistent with a previous report[20]. Genetic ablation of MTX1, MTX2 or MTX3 did not affect the migration of the MICOS complex **(Fig. 1D-F)**. While the loss of MTX1 or MTX3 had no effect on the migration of the SAM complex, we found that the SAM complex migrated faster in MTX2^KO^ mitochondria when compared to control mitochondria, an effect that was corrected by complementation with ^FLAG^MTX2 **(Fig. 1E)**. Collectively, we demonstrate that although MTX2 is required for the stability of MTX1 and the SAM complex, the loss of MTX1 or MTX3 does not affect the stability of this complex or the levels of MTX2.

**Fig. 1.**
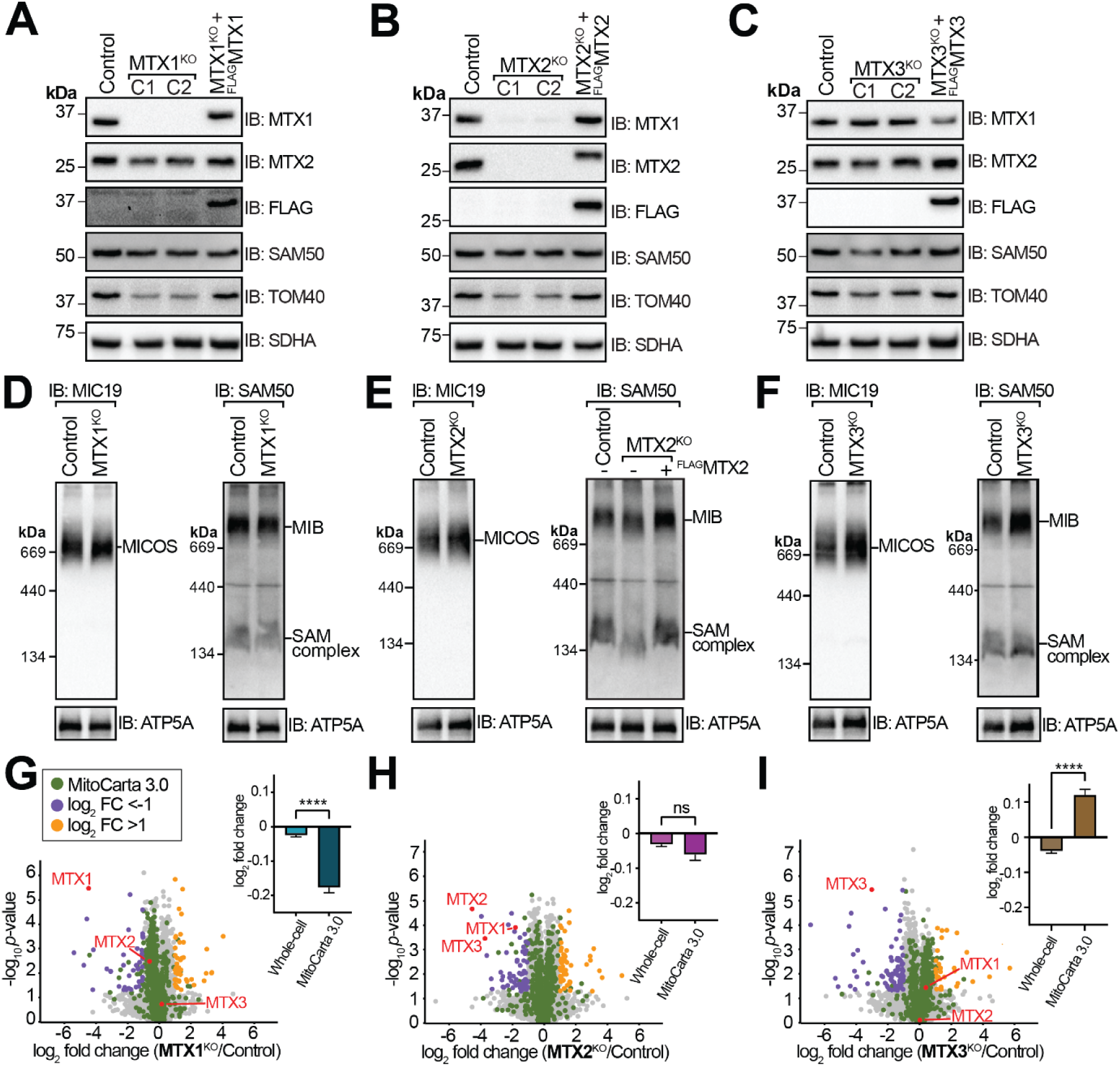
Loss of MTX1 or MTX3 differentially alters mitochondrial protein content. **A-C)** Mitochondria isolated from control, Metaxin KO clones (C1 and C2) and rescue cells were subjected to SDS-PAGE or **D-F)** BN-PAGE and analysis by immunoblotting. Loading controls were SDHA or ATP5A as indicated. **G-I)** Volcano plots illustrating changes to global cellular protein abundance in response to the loss of **G)** MTX1, **H)** MTX2 or **I)** MTX3. Proteins were considered significantly changed where the log_2_ fold change was >1 or <-1 and *p*-value <0.05 (two-sided students’ *t*-test). N=3 biological replicates. For each data set, the mean log_2_ fold change was (±SEM) of all detected proteins and all detected MitoCarta 3.0 proteins was plotted. Statistical comparisons were performed using a one-sided Student’s *t*-test (**** *p*<0.0001).

Next, we explored the response of the global cellular proteome to MTX1, MTX2 or MTX3 loss using liquid chromatography-tandem mass spectrometry (LC/MS-MS) **(Fig. 1G-I, Table S1)**. In the absence of MTX1, we observed a significant reduction in mitochondrial protein abundance relative to whole-cell protein content **(Fig. 1G)**. In addition, the levels of MTX3 were unchanged in response to MTX1 loss (**Fig. 1G)**. The loss of MTX3 did not significantly affect the levels of MTX1 or MTX2, however, we observed a significant increase in mitochondrial protein abundance in MTX3^KO^ cells, suggesting that MTX1 and MTX3 function independently in the regulation of mitochondrial protein content **(Fig. 1I)**. Our analysis of the MTX2^KO^ proteome revealed that MTX2 is required for the stability of both MTX1 and MTX3, suggesting that a clear hierarchy of stability exists among the subunits of the human SAM complex **(Fig. 1H)**. Despite this, MTX2 loss did not lead to significant changes in the whole-mitochondrial proteome **(Fig. 1H)**. Functional enrichment analysis was performed by assigning Gene Ontology (GO) terms to significantly altered proteins[21] **(Fig. S2A and S2B)**. Proteins associated with mitochondrial translation and protein targeting to the mitochondrion, including subunits of the TOM and translocase of the inner membrane (TIM) complexes, were significantly decreased in MTX1^KO^ cells **(Fig. S2A and Table S2)**. Meanwhile, proteins associated with actin cytoskeleton organization were significantly decreased in cells lacking MTX2 **(Fig. S2A and Table S2)**. We found pathways that were exclusively enriched among significantly increased proteins in MTX3^KO^ cells which included OXPHOS and ATP synthesis **(Fig. S2B and Table S3)**. Collectively, our proteomic analysis allowed us to uncover unique pathways that are exclusively perturbed in the absence of MTX1 or MTX3, suggesting that each protein makes discrete contributions to mitochondrial health.

### β-barrel biogenesis is impaired in response to the loss of Metaxins

Previous work demonstrated that MTX2 depletion in human cells impairs VDAC1 biogenesis *in vitro*[19]. To date, the requirement of MTX1 and MTX3 for the biogenesis of β-barrel proteins in human mitochondria has not been shown definitively. Our proteomic characterization revealed that the steady-state abundance of some β-barrel proteins was reduced in MTX1^KO^ and MTX2^KO^ cells but not MTX3^KO^ cells **(Fig. S2C-E)**. To resolve whether MTX1 and MTX3 were required for β-barrel biogenesis, we assayed the import of human SAM complex substrates, VDAC1 and TOM40, into mitochondria isolated from MTX1^KO^, MTX2^KO^ and MTX3^KO^ cells using [^35^S]-labelled β-barrel precursor proteins[19,20]. As expected[19], we observed a significant reduction in VDAC1 import and assembly in mitochondria isolated from MTX2^KO^ cells compared to control mitochondria **(Fig. 2A and 2C)**. Similarly, VDAC1 biogenesis was significantly impaired in MTX1^KO^ mitochondria. However, no VDAC1 import and assembly defects were observed in MTX3^KO^ mitochondria **(Fig. 2A and 2C)**. We also uncovered a reduction in steady-state VDAC1 levels in MTX1^KO^ mitochondria indicating that VDAC deficiency occurs independent of loss of total cellular mitochondrial content in MTX1^KO^ cells **(Fig. 2A)**. Collectively, we uncovered a previously unappreciated distinction in the requirement of MTX1 and MTX3 for the import and assembly of VDAC1.

**Fig. 2.**
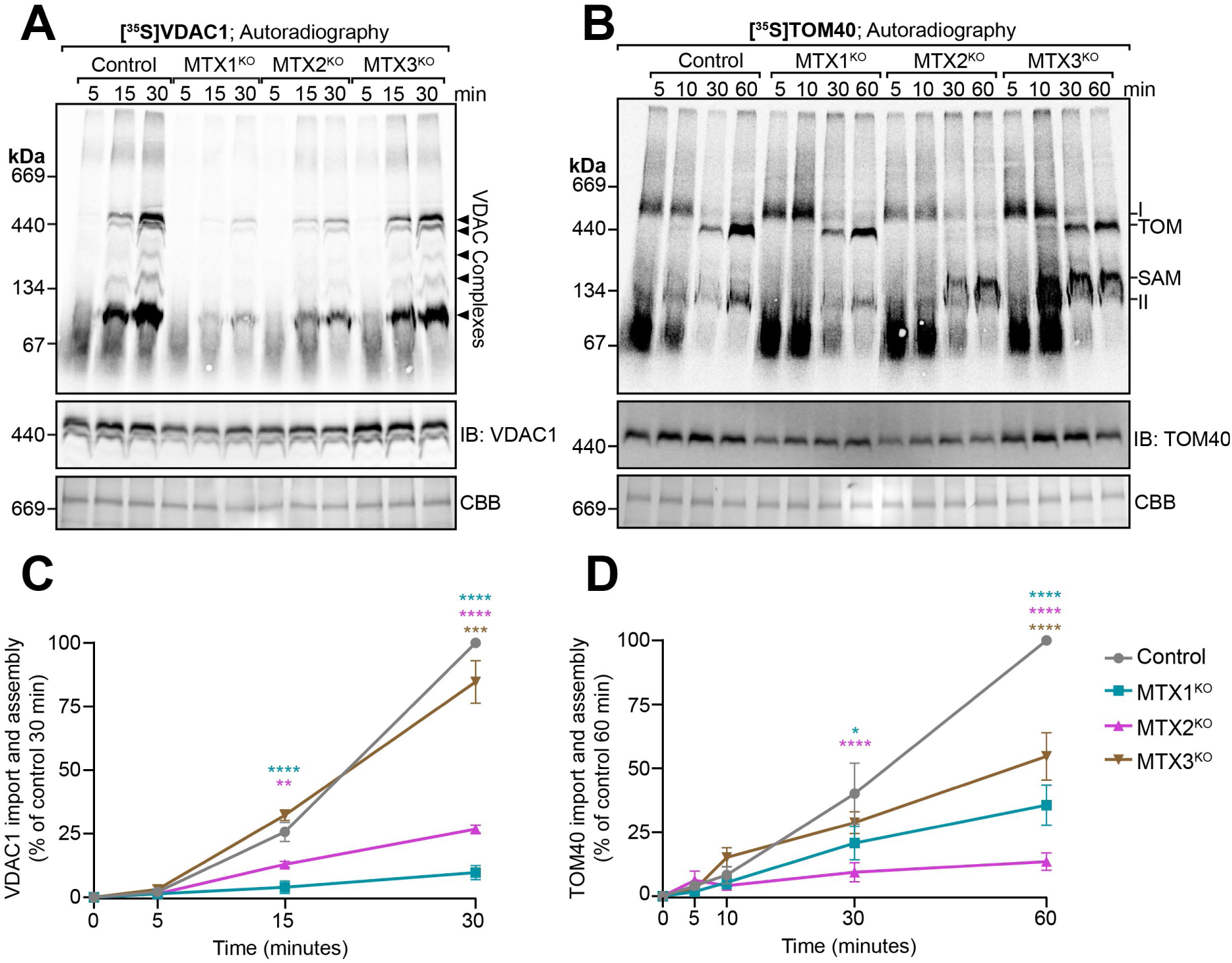
Loss of Metaxins differentially affects β-barrel biogenesis. *In vitro* translated **A)** [^35^S]VDAC1 or **B)** [^35^S]TOM40 was imported into mitochondria isolated from control, MTX1^KO^, MTX2^KO^, and MTX3^KO^ cells for times as indicated. Mitochondria were solubilized in 1% digitonin and the resulting complexes were resolved by BN-PAGE and imaged by phosphorimaging. Immunoblotting of steady-state complexes was performed using antibodies against VDAC1 or TOM40. Coomassie Brilliant Blue (CBB) staining of the PVDF was used as a loading control. **C)** Mature [^35^S]VDAC1 (two ∼440kDa species) or **D)** [^35^S]TOM40 complexes (TOM) were quantified by densitometry at each time point and expressed as a percentage against the terminal control time point. Statistically significant differences between genotypes at each time point were determined using a two-way ANOVA with Dunnett’s multiple comparisons test (n=3 experimental replicates, * *p*<0.05, ** *p*<0.01, *** *p*<0.001, **** *p*<0.0001). Data presented as mean ± SEM.

TOM40 import and assembly is captured in three distinct assembly states - precursor TOM40 bound to the endogenous TOM complex before translocating the OMM (Assembly I), the newly-folded TOM40 β-barrel associated with small TOM proteins (Assembly II), and its integration into the mature TOM complex (TOM)[20]. In the absence of MTX1, TOM40 followed the canonical import and assembly pathway described above, however its assembly kinetics into the mature TOM complex were reduced in comparison to control mitochondria **(Fig. 2B and 2D)**. In mitochondria isolated from both MTX2^KO^ and MTX3^KO^ cells we also observed newly-imported TOM40 accumulating in an additional intermediate consistent with the SAM complex (SAM)[20] **(Fig. 2B)**. Transfer of TOM40 from the SAM-TOM intermediate to the mature TOM complex was significantly impaired in MTX2^KO^ cells, but less so in MTX3^KO^ cells, suggesting that MTX2 may be required for the later stages of TOM40 maturation. Taken together, these findings suggest that the import and assembly pathway of TOM40 is altered in the absence of MTX2 and MTX3. Meanwhile, loss of MTX1 showed reduced efficiency of TOM40 import and assembly.

### Reduced mitochondrial volume and network collapse are driven by ablation of MTX1

Given that we observed opposing changes in mitochondrial protein content in MTX1^KO^ and MTX3^KO^ cells, we investigated whether mitochondrial volume, morphology, or distribution of mtDNA were affected in these cell lines. Using immunofluorescence microscopy, we found that MTX1 loss caused network-wide collapse of the mitochondrial reticulum characterized by swollen and misshapen mitochondria with reduced interconnectivity **(Fig. 3A)**. Furthermore, we identified subpopulations of mitochondria which contained very few mtDNA puncta relative to their size and others which were enlarged but contained several mtDNA puncta with irregular distribution **(Fig. 3A)**. MTX2^KO^ mitochondria appeared swollen, however, we did not observe any changes to mtDNA distribution **(Fig. 3A)**. By contrast, MTX3^KO^ mitochondria maintained their interconnectedness, and were noticeably thinner than those of control cells **(Fig. 3A)**. Similar to MTX1^KO^ cells, mtDNA distribution was altered in the absence of MTX3, with some mitochondria, particularly those more distal to the perinuclear area, containing very few mtDNA puncta **(Fig. 3A)**. Although mtDNA distribution was altered in MTX1^KO^ and MTX3^KO^ cells, we did not observe any changes to total mtDNA content across all three genotypes **(Fig. S3B)**.

**Fig. 3.**
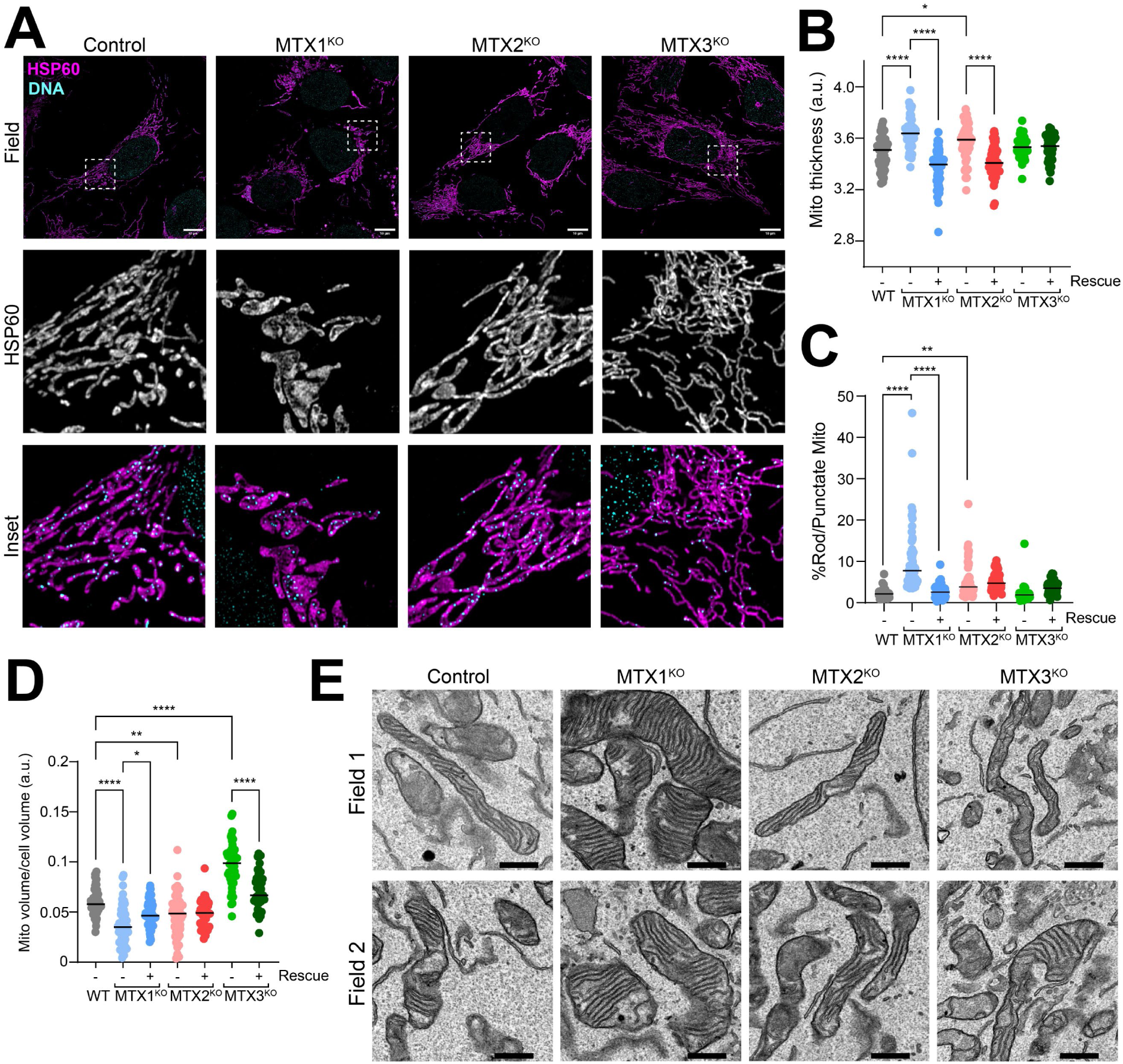
Mitochondrial morphology and ultrastructure are altered in the absence of MTX1, MTX2 or MTX3. **A)** Representative confocal images of mitochondrial morphology (antibodies against HSP60) and mtDNA content (antibodies against DNA) in control, MTX1^KO^, MTX2^KO^, and MTX3^KO^ cells. Inset represents optical zoom of area indicated by hatched box imaging for HSP60 (white) or merged to show co-localization with DNA. Scale bar = 10µm. Mitochondria were quantified for **B)** thickness **C)** morphology (% of mitochondria classified as “rod” or “punctate”) and **D)** total volume according to NDUFAF2 staining in control (n=50 cells), MTX1^KO^ (n=56 cells), MTX1^KO^ + ^FLAG^MTX1 (n=52 cells), MTX2^KO^ (n=47 cells), MTX2^KO^ + ^FLAG^MTX2 (n=50 cells), MTX3^KO^ (n=47 cells) and MTX3^KO^ + ^FLAG^MTX3 (n=47 cells) cells. All statistical tests were performed using an ordinary one-way ANOVA with Tukey’s multiple comparisons test (ns *p*>0.05, * *p*<0.05, ** *p*<0.01, \*\*\*\**p*<0.0001). **E)** Representative TEM micrographs of control, MTX1^KO^, MTX2^KO^ and MTX3^KO^ cells. Scale bar = 500nm.

Next, we complemented our qualitative findings with volumetric image analysis using antibodies corresponding to NDUFAF2 as a mitochondrial matrix marker **(Fig. S3A, C-E)**. Since MTX2 deficiency has been reported to cause abnormal nuclear architecture[14], we also used antibodies against the nuclear envelope protein Lamin A/C to examine whether loss of MTX1 or MTX3 was also linked to changes in nuclear morphology **(Fig. S3A)**. Loss of MTX1, and to a lesser extent MTX2, was linked to an increase in network-wide mitochondrial swelling, which was corrected upon complementation with ^FLAG^MTX1 or ^FLAG^MTX2 respectively **(Fig. 3B, Fig. S3A)**. Genetic ablation of MTX1 was also associated with a significant increase in the proportion of mitochondria classified as “rod” or “punctate” in appearance in tandem with a reduction in those classified as “branched” or “elongated”, indicating mitochondrial interconnectivity **(Fig. 3C and S3C)**. Consistent with our proteomic data, we observed a significant decrease in mitochondrial volume in MTX1^KO^ cells **(Fig. 3D)**. Inversely, we found the mitochondrial volume of MTX3^KO^ cells to be significantly increased, also consistent with our proteomic data **(Fig. 3D)**. In both instances, these effects were corrected following re-expression of ^FLAG^MTX1 and ^FLAG^MTX3 respectively. Curiously, we observed a significant decrease in mitochondrial volume in MTX2^KO^ cells, albeit to a lesser extent than MTX1^KO^ cells and with greater population-wide variability **(Fig. 3D)**. We also observed an increase in total mitochondrial volume occupying the perinuclear region in MTX1^KO^ cells although the effect was minor **(Fig. S3D)**. Contrary to a previous report[14], our analysis of nuclear morphology did not uncover any significant differences in nuclear irregularity in any of our KO cell lines **(Fig. S3A and S3E)**. Next, we assessed mitochondrial ultrastructure using Transmission Electron Microscopy (TEM) **(Fig. 3E)**. Although the loss of MTX1 did not have any obvious effect on cristae architecture, the enlarged mitochondria observed in this genotype appeared more cristae-dense than those of control cells **(Fig. 3E)**. Consistent with a recent report[22], we observed some mitochondria with wider cristae in cells lacking MTX3 than those of control cells **(Fig. 3E)**. Collectively, our analysis revealed clear alterations to mitochondrial morphology, distribution and volume driven by the loss of MTX1 and showed that genetic ablation of MTX3 produces a distinct phenotype to that of MTX1^KO^ cells.

### MTX1 and MTX3 exist in unique assembly states and have mutually exclusive interacting partners

According to a previous report, MTX2 and MTX3 co-migrate with a SAM50-containing complex of approximately 150kDa in size, whereas MTX1 co-migrates with the OMM MIB-associated protein, DNAJC11[23], in a species of approximately 250kDa and 450kDa[13]. The responsiveness of these assembly states to the loss of any one of the Metaxin proteins has not been investigated. Analysis of MTX1- and MTX3-containing complexes by 1D BN-PAGE has been constrained by the unavailability of commercial antibodies that can detect MTX1 or MTX3 within their native assembly states. To overcome this obstacle, we undertook *in vitro* import and assembly analysis of [^35^S]-labelled MTX1, MTX2 or MTX3 precursor protein. We first compared the assembly profiles of MTX1, MTX2 and MTX3 in control mitochondria **(Fig. 4A)**. Akin to previous reports[13,19,24], MTX1 was primarily detected in a species of approximately 50-70kDa while a species of lower abundance was observed at 400kDa, although the signal acquired from [^35^S]MTX1 import was relatively weak **(Fig. 4A)**. MTX2 was detected in a species of approximately 150kDa as well as a very high molecular weight species which correlated with a sub-assembly of the MIB complex **(Fig. 4A)**[13]. MTX3 was detected in the same high molecular weight species as MTX2, however, we also identified an additional, unique assembly state of approximately 250kDa **(Fig. 4A)**. The import and assembly of MTX1 and MTX2 was unaffected across all genotypes assessed, although assembly appeared more efficient when imported into mitochondria lacking the endogenous protein **(Fig. 4B-C)**. In contrast, the import and assembly of MTX3 into both assembly states identified was significantly impaired in mitochondria lacking MTX2 and was comparable to control levels in mitochondria lacking endogenous MTX3 **(Fig. 4B-D)**. Collectively, we reveal a previously unreported dependency of MTX3 on MTX2 for its capacity to assemble into its native complexes. Furthermore, we found that MTX1 does not depend on MTX2 for its import and assembly, despite MTX2 being required for the maintenance of MTX1 protein levels at steady state.

**Fig. 4.**
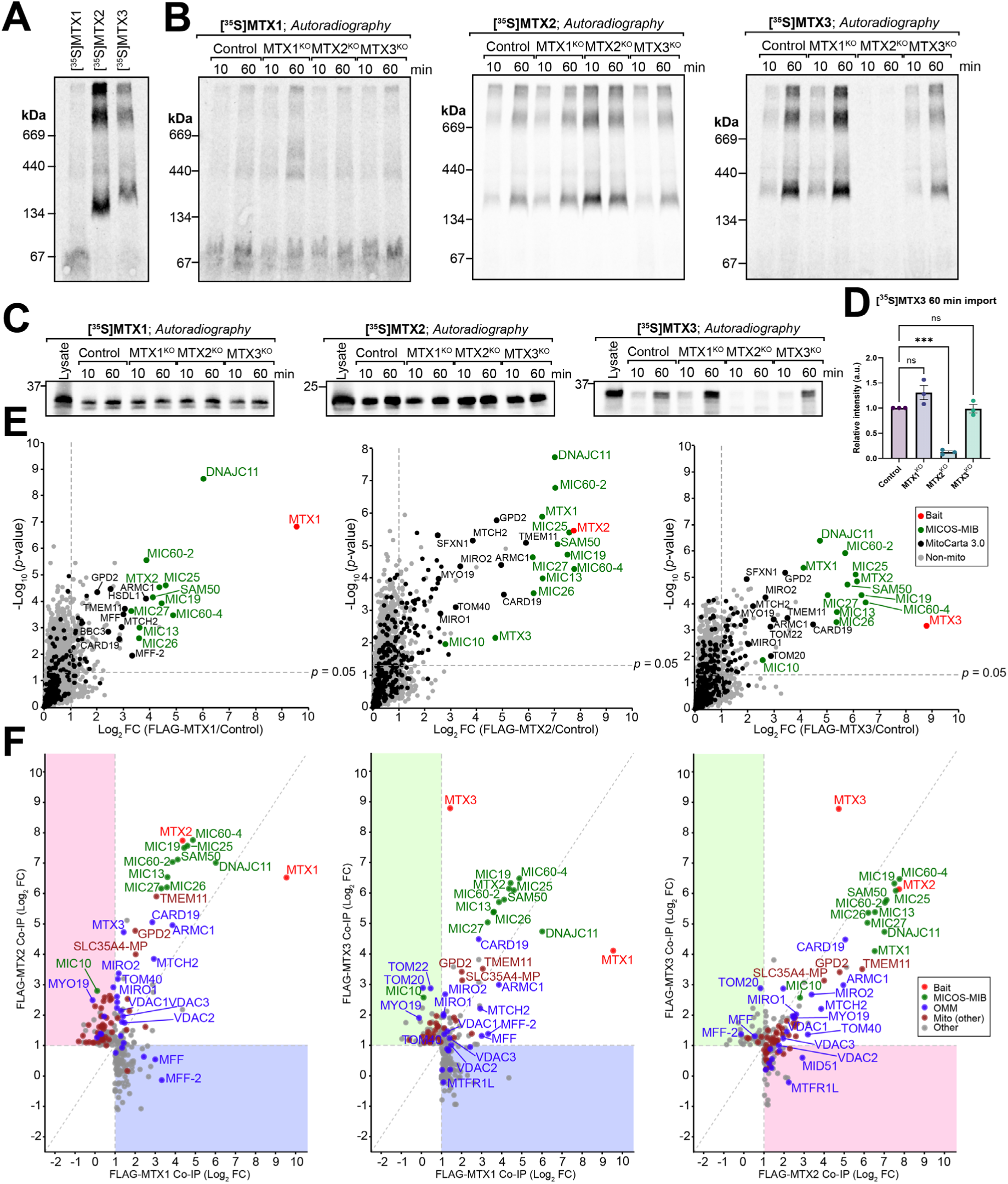
MTX1 and MTX3 have discrete assembly states and unique interacting partners. **A)** [^35^S]MTX1, [^35^S]MTX2 and [^35^S]MTX3 were imported into mitochondria isolated from control cells for 60 minutes at 37°C and analysed by BN-PAGE and phosphor imaging. **B-C)** [^35^S]MTX1, [^35^S]MTX2 and [^35^S]MTX3 were imported into mitochondria isolated from control, MTX1^KO^, MTX2^KO^ and MTX3^KO^ cells for 10 and 60 minutes at 37°C and were analysed by **B)** BN-PAGE or **C)** SDS-PAGE followed by phosphor imaging. **D)** The amount of [^35^S]MTX3 imported into mitochondria isolated from control, MTX1^KO^, MTX2^KO^ and MTX3^KO^ cells after 60 minutes was quantified and statistical significance was determined using an ordinary one-way ANOVA with Dunnett’s multiple comparison’s test (*** *p*<0.001, n=3 experimental replicates). **E)** Volcano plots illustrating the interactomes of ^FLAG^MTX1, ^FLAG^MTX2 and ^FLAG^MTX3 (left to right). Significantly enriched proteins were identified following log_2_ transformation of the data followed by Student’s *t*-test (*p*<0.05, indicated by horizontal dotted line). Vertical dotted line indicates manually asserted fold change significance cut off (FC>1). **F)** Fold change values of proteins with FC>1 and *p*>0.05 from at least one of two interactomes compared were plotted to identify unique interactors of each proteome. Shaded areas indicate regions where unique interactors are plotted (blue = unique to MTX1, pink = unique to MTX2, green = unique to MTX3).

We next investigated whether MTX1 and MTX3 have unique interacting partners given their discrete assembly profiles. Using affinity enrichment mass spectrometry (AE-MS), we defined the steady-state interactomes of MTX1, MTX2 and MTX3 **(Fig. 4E, Table S4)**. Known MICOS-MIB proteins were among the most highly enriched interactors of all three Metaxin proteins **(Fig. 4E)**. TMEM11, ARMC1 and CARD19, which have been highlighted in previous reports as associates of the MICOS-MIB complex [25–28], were also strongly enriched across all three interactomes **(Fig. 4E)**. To discern the unique interacting proteins of MTX1, MTX2 and MTX3, we compared proteins that were significantly enriched in one interactome but not in another **(Fig. 4F)**. We noted that two isoforms of Mitochondrial Fission Factor (MFF), an adaptor for the large GTPase, DRP1[29,30], were significantly enriched in both the MTX1 and MTX3 interactomes, but not the MTX2 interactome **(Fig. 4F)**. Similarly, we found that another regulator of mitochondrial dynamics, MTFR1L[31], was present in the MTX1 and MTX2 interactome but not MTX3 **(Fig. 4F)**. Metaxins have been previously implicated in the formation of complexes with mitochondrial trafficking adaptors that facilitate mitochondrial motility along microtubules[32,33]. Furthermore, the unconventional myosin, MYO19, which is required for linking mitochondria to the actin cytoskeleton, has been previously described as an interactor of MTX3[22,34]. We reaffirmed this finding in our own data and further uncovered that MTX2, but not MTX1, is a stable interactor of MYO19. The mitochondrial Rho GTPases MIRO1 and MIRO2, which are adaptors not only for MYO19[35] but for mitochondrial trafficking adaptors TRAK1 and TRAK2[36], were significantly enriched across all three proteomes **(Fig. 4E and 4F)**. We did not detect cytosolic trafficking adaptors, TRAK1 and TRAK2, in any of the Metaxin interactomes, however, we speculate that this is due to the transient nature of this interaction. Given the differences in lower-order assembly states occupied by MTX1, MTX2 and MTX3, we sought to discern interactors that could account for the size differences in these assembly states. One previous study has demonstrated that MTX1 co-migrates with a complex containing TOM40, TOM22 and TOM20[24]. Contrary to this study, we found that neither TOM20 nor TOM22 were significantly enriched with ^FLAG^MTX1 but rather were unique interactors of MTX3 when comparing the MTX1 and MTX3 interactomes **(Fig. 4F)**. Furthermore, we were able to discern TOM20, but not TOM22, as an interactor of MTX3 but not MTX2. To conclude, we discovered unique interactors of the Metaxin proteins in support of their existence in distinct assembly states.

## Discussion

In higher-order eukaryotes, the mitochondrial proteome has become progressively more diverse due to the duplication and diversification of nuclear-encoded mitochondrial genes[37,38]. MTX1 and MTX3 are examples of human mitochondrial proteins that arose from gene duplication, yet the selective advantage of two homologous SAM complex accessory subunits in humans, and whether they have diverged in function, has not been explored. Our analysis of cells lacking the human Metaxin proteins revealed clear differences in the biogenesis of SAM complex substrates and in maintaining the mitochondrial proteome. Indeed, our analysis of MTX1 and MTX3 demonstrated that these proteins exist in unique molecular weight assembly states, as previously suggested[13]. Unexpectedly, we found that while MTX1 could assemble into higher-molecular-weight complexes independently of the other Metaxin proteins, as MTX3 import and assembly was exclusively dependent on MTX2. Given that the loss of either MTX1 or MTX3 appears to have opposing effects on the mitochondrial proteome, the diversification of the homologs could be a way to fine-tune the import and biogenesis of mitochondrial proteins, acting as a rheostat to optimize mitochondrial function. Taken together, our data suggest that MTX1 and MTX3 exist in distinct SAM50-containing complexes that have unique roles in mitochondrial protein biogenesis.

Before folding and insertion into the OMM, β-barrel substrates must be imported into the IMS via the TOM complex and then chaperoned to the IMS-face of the SAM complex with the aid of small TIM proteins[39]. Together with our finding that β-barrel import is perturbed in the absence of MTX1, our proteomic analysis of MTX1^KO^ cells uncovered a reduction in proteins associated with both the TOM and TIM complexes. Defective protein import into MTX1^KO^ mitochondria would account for both the loss of overall mitochondrial protein content as well as the perturbed capacity of MTX1^KO^ mitochondria to facilitate the biogenesis of SAM complex substrates. However, since assembly of TOM40 requires the SAM complex[8], distinguishing a SAM complex defect from a MTX1-specific role in β-barrel biogenesis in the absence of MTX1 is challenging. The yeast MTX2 homolog, SAM35, is required for recognising the conserved β-signal of SAM complex substrates in the early stages of β-barrel maturation, with a mutant SAM35 leading to defective TOM40 assembly into the mature TOM complex[40]. Similarly, our data shows that in the absence of MTX2, TOM40 is unable to effectively integrate into the mature TOM complex, however, it does not affect the capacity of precursor TOM40 to associate with the endogenous TOM complex or the SAM complex. Since the levels of MTX1 and MTX3 are strongly reduced in MTX2^KO^ mitochondria, this suggests that SAM50 alone maintains some capacity for β-barrel integration.

Analysis of proteomic changes resulting from loss of Metaxins revealed that MTX2 does not depend on MTX1 or MTX3 for stability, whereas loss of MTX2 caused severe depletion of both MTX1 and MTX3. Unlike the yeast SAM complex[11,12,41], the structure of the human SAM complex has not been solved, and so how the subunits associate with each other and the OMM is not fully understood. According to the structure of the yeast SAM^core^ complex, SAM37 does not directly contact SAM50 on the cytosolic face of the complex[11]. Rather SAM37 associates with SAM50 indirectly via interactions between SAM35 on the cytosolic face and directly with SAM50 through an N-terminal segment that reaches across the OMM and into the IMS[11]. Based on their likely topologies[42,43], we anticipate that MTX1 and MTX3 interact with MTX2 in a similar manner to SAM37 and SAM35. Since MTX2 is stable in the absence of MTX1 or MTX3, we would expect that the arrangement of MTX1/3 and MTX2 in the human SAM or MIB complex would resemble that of the yeast SAM complex.

Interestingly, MTX1 was recently described as a component of an ARMC1-containing complex that also contained DNAJC11 and MTCH2[27]. Our AE-MS analysis confirmed that MTX1 strongly associates with DNAJC11, as reported previously[44], as well as ARMC1 and MTCH2. Each of these proteins is an interactor of known MIB complex components[25,45,46]. While MTX2 was not found to be part of the ARMC1-DNAJC11-MTX1-MTCH2 complex in the prior study[27], our analysis of MTX2 interacting proteins identified all of these components, suggesting that this complex may also interact with the SAM complex. Indeed, since MTX2 is largely required for MTX1 stability, future work to understand how MTX1 partitions between the SAM/MIB and the ARMC1-DNAJC11-MTX1-MTCH2 complex may uncover further aspects of mitochondrial distribution and protein biogenesis.

The mitochondrial phenotype associated with MTX1 loss was characterized by loss of mitochondrial interconnectivity, swelling and abnormal inter-mitochondrial mtDNA distribution. This phenotype is in contrast with that of MTX3^KO^ cells which have increased mitochondrial interconnectivity but otherwise resembled control mitochondria. Moreover, MTX1 loss caused a decrease in total mitochondrial volume whereas mitochondrial volume was elevated in MTX3^KO^ cells. Previous findings using an siRNA-mediated knockdown of MTX1 or MTX3 depletion, found that reduced MTX1 caused an increase in mitochondrial content, while MTX3 depletion did not affect mitochondrial content[47]. We speculate that knockdown may not cause sufficient MTX3 protein loss to elicit a noticeable effect on mitochondrial protein content. In contrast, the model of MTX3 loss described in our work is stable and complete rather than transient. Mitochondrial swelling is typically associated with opening of the Mitochondrial Permeability Transition Pore (mPTP) on the IMM, which causes the matrix to swell leading to IMM morphology changes and a loss of cristae organization[48,49]. Our TEM analysis of enlarged mitochondria in MTX1^KO^ cells did not reveal any changes to IMM architecture that resembled mPTP-induced matrix swelling, suggesting that another mechanism may be responsible for changes in MTX1^KO^ mitochondrial morphology. Enlarged mitochondria are often the result of defective mitochondrial clearance, metabolic stress or impaired mitochondrial dynamics[50–52]. Our data indicates that the OMM fission adaptor MFF, is preferentially enriched with MTX1. MFF is required for the recruitment of DRP1 to the surface of mitochondria fated for division[29,30]. Cells deficient for the fission executor DRP1 exhibit enlarged mitochondria throughout the network which contain densely stacked cristae that resemble those observed in MTX1^KO^ cells[53]. MFF participates in midzone fission which occurs near the centre of mitochondria and promotes mitochondrial biogenesis[54]. Dysfunctional midzone fission in MTX1^KO^ cells may explain why MTX1^KO^ cells lose overall mitochondrial mass without loss of total mtDNA.

Mutations in the *MTX2* gene manifest as a rare progeroid syndrome in patients known as MADaM[14,15]. Our characterization of MTX2^KO^ cells uncovered that both MTX1 and, to a greater extent, MTX3, are significantly depleted upon loss of MTX2, highlighting the importance of understanding the effect of MTX1 and MTX3 loss in isolation. MADaM patient-derived fibroblasts show altered mitochondrial morphology, which has been described as mitochondria having reduced interconnectivity and a loss of individual mitochondria[14]. Indeed, from our findings, MTX1 loss is sufficient to cause mitochondrial network collapse and reduced mitochondrial content. Moreover, we uncovered a defect in TOM40 biogenesis in MTX2^KO^ and, to a lesser extent, in MTX1^KO^ mitochondria. The contribution of defective mitochondrial protein import to the pathogenesis of MADaM is a compelling direction for future investigation into understanding the molecular mechanisms contributing to this disease. Curiously, mutations in the *TOM7* gene, encoding a component of the TOM complex, are linked to the progeroid syndrome known as Garg-Mishra Progeroid Syndrome (GMPGS)[55,56]. The disease-causing TOM7^Trp25Arg^ mutation is predicted to interfere with the association between TOM7 and TOM40, suggesting that defective protein import may be responsible for the manifestation of GMPGS[56]. Although it is possible that defects in mitochondrial protein import lead to a progeria phenotype, this remains to be firmly established. Given that we also did not observe changes in nuclear morphology in MTX2^KO^ cells, progeroid-like symptoms in MADaM patients may be due to underlying defects in mitochondrial protein import. Collectively, our findings provide novel insights into the possible contributions of MTX1 and MTX3 depletion to the phenotype observed in patients with null mutations in the *MTX2* gene.

## Methods and Materials

### Plasmids and molecular cloning

Targeted gene disruption for knockout cell line generation was achieved by selecting two gene-specific guide RNAs (gRNAs) using CHOPCHOP[57]. Details of gRNA sequences are listed in **Table S5**. Hybridized gRNA-specific oligonucleotides were cloned into BbsI-digested pSp-Cas9(BB)-2A-GFP (a gift from F. Zhang; Addgene, PX458) or pSp-Cas9(BB)-2A-mCherry (generated as part of this study). pSp-Cas9(BB)-2A-mCherry was generated by restriction enzyme excision of the GFP cDNA using EcoRI and ligation of mCherry cDNA PCR-amplified from a plasmid template using primers designed to introduce a G>A mutation at position 432 to remove the mCherry BbsI RE site. Inserts for cloning into pBMN-Z (Addgene #1734) or pBABE-puro (Addgene #1764) for stable protein expression were amplified from plasmid templates by PCR. Forward primers were designed to incorporate a FLAG^®^ epitope tag (5′-GACTACAAAGACGATGACGACAAG-3′) at the 5’ end of the cDNA of interest. All plasmids cloned as part of this study were sequence verified by Micromon Genomics (Monash University).

### Cell culture

Human bone osteosarcoma (U2OS) cells were cultured in Roswell Park Memorial Institute (RPMI) medium supplemented with 10% foetal bovine serum (FBS), 1% penicillin-streptomycin (P/S, Sigma-Aldrich, P4333), 1× GlutaMAX^TM^ (Thermo Fisher Scientific, 35050061) and 50µg/mL uridine (MP Biomedicals, 0219476350). Cells were continually cultured at 37°C in a humidified 5% CO_2_ incubator.

### Transfection and stable cell line generation

Transient transfection of cultured cells was performed using Lipofectamine LTX^TM^ with PLUS^TM^ reagent according to manufacturer’s instructions. Generation of pantropic retrovirus for stable protein expression was achieved by co-transfecting HEK293T cells with the transfer plasmid encoding cDNA of interest in combination with retroviral helper vectors, Gag-Pol (Addgene #14887) and VSV-G. Viral supernatants were collected 48 hours post-transfection and transferred to target cells with the addition of 8µg/mL polybrene. Cells were selected for stable protein expression by culturing in 1µg/mL puromycin for a minimum of 48 hours. Following puromycin selection, clonal populations of cells were derived from the mixed cell population using fluorescence-activated cell sorting (FACS) and were screened for protein expression by SDS-PAGE and immunoblotting.

### Gene editing and screening

CRISPR/Cas9-mediated genome editing was performed as described previously[58]. pSp-Cas9(BB)-2A-GFP (PX458) and pSp-Cas9(BB)-2A-mCherry plasmids encoding target-specific oligonucleotide sequences were transiently co-transfected into wild-type (WT) U2OS cells and selected by FACS for high expression of GFP and mCherry. Clonal cell populations were screened by extracting genomic DNA (gDNA) using QuickExtract^TM^ DNA Extraction solution (LGC Biosearch Technologies, QE09050) and genotyped using primers flanking the expected dropout region as well as an “internal” primer designed to bind within the excision region. Genotyping primers are listed in **Table S6**. Where antibodies against the target protein were available, protein loss was confirmed by SDS-PAGE and immunoblotting.

### Mitochondrial isolation

Crude mitochondrial extracts were prepared using methods described previously[59,60]. The resulting mitochondrial preparation was resuspended in sucrose storage buffer (10 mM HEPES pH 7.6, 0.5 M sucrose) and stored at −80°C. Protein content of the resulting preparation was determined using a bicinchoninic acid (BCA) assay kit (Thermo Fisher Scientific, 23227).

### Gel electrophoresis, western transfer and immunoblotting

Protein separation using a tris-tricine SDS-PAGE system or Blue Native (BN)-PAGE was performed using methods described previously[59,61,62]. Samples were run alongside Bio-Rad Precision Plus Protein^TM^ Kaleidoscope^TM^ Prestained Protein Standards (Bio-Rad, 1610375). Following electrophoresis, proteins were transferred onto a PVDF membrane (Merck, IPVH00010) using either a Power Blotter XL System (Invitrogen, PB0013) or a Novex^TM^ Semi-Dry Horizontal Blotter (Invitrogen, SD1000) according to manufacturers’ directions. Primary antibodies used for immunoblotting and relevant dilutions are listed in **Table S7**. Membranes were incubated in primary antibodies overnight at 4°C. Horse radish peroxidase-conjugated anti-mouse (Sigma-Aldrich, A9044) or anti-rabbit (Sigma-Aldrich, A0454) secondary antibodies were prepared as a 1:10,000 dilution in 5% w/v skim milk in TBS-T. Immunoblots were developed using Clarity^TM^ Western Electrochemiluminescence Substrate (Bio-Rad, 1705061). Images were acquired using a Chemidoc^TM^ XRS+ Imaging System (Bio-Rad, 1708265).

### *In vitro* mitochondrial protein import

*In vitro*-coupled transcription and translation of [^35^S]-labelled precursor proteins was performed using the TnT® T7/SP6 Quick Coupled Transcription/Translation System (Promega, L1170/L2080) according to manufacturer’s directions. Templates for use with the TnT® T7 Quick Coupled Transcription/Translation System were PCR-amplified from a plasmid template with a forward primer designed to incorporate a T7 promotor (5’-TAATACGACTCACTATAAGGGAGA-3’) directly upstream of the open reading frame (ORF). For *in vitro* transcription/translation of [^35^S]TOM40, pGEM-4z encoding TOM40 cDNA downstream of the SP6 promotor was used as the template[20]. Mitochondrial protein import was performed according to procedures described previously[59,60]. Radiolabelled proteins were incubated with isolated mitochondria suspended in import buffer (250 mM sucrose, 5 mM magnesium acetate, 80 mM potassium acetate, 5 mM ATP, 10 mM sodium succinate, 1 mM DTT, 20 mM HEPES-KOH pH 7.4). Import of [^35^S]VDAC1 was performed at 25°C. All other precursors were imported at 37°C. Following import, isolated mitochondria were prepared for BN-PAGE or SDS-PAGE and results were visualized by phosphorimaging.

### Affinity enrichment of epitope-tagged proteins by co-immunoprecipitation (Co-IP)

Co-IP was performed to identify protein-protein interactions as described previously with some modifications[59]. Whole-cell protein was solubilized using 1% w/v digitonin prepared in solubilization buffer (20 mM Tris pH 7.4, 50 mM NaCl, 10% v/v glycerol, 0.1 mM EDTA) supplemented with 0.125U/mL Turbonuclease from *Serratia marcescens* (Sigma-Aldrich, T4330) and cOmplete^TM^, EDTA-free Protease Inhibitor Cocktail (Roche, 11873580001). Samples were incubated on ice for 30 minutes and clarified by centrifugation at 16,000 *g* for 10 minutes at 4°C. Anti-FLAG^®^ M2 Affinity Gel beads (Sigma-Aldrich, A2220) were equilibrated by resuspending and washing in solubilization buffer (detergent and nuclease omitted) followed by wash buffer (solubilization buffer supplemented with 0.2% w/v digitonin). Equal volumes of beads were transferred to Pierce^TM^ Screw Cap Spin Columns (Thermo Fisher Scientific, 69705). Clarified whole-cell lysates were added to spin columns with beads and incubated on an end-over-end rotator for 2 hours at 4°C. Beads were washed 5 times with 0.2% w/v digitonin in solubilization buffer on end-over-end rotator and proteins were eluted using 150 µg/mL DYKDDDDK tag peptide (Assay Matrix, A6002). Precipitation of eluted protein was achieved with the addition of ice-cold acetone to each sample followed by overnight incubation at −20°C. Precipitated material was dried either in a fume cupboard or by vacuum centrifugation.

### Sample preparation for mass spectrometry

Peptides for analysis by mass spectrometry were prepared using S-Trap^TM^ Mini Columns (ProtiFi, CO2-mini-80) according to manufacturer’s instructions. On-column tryptic digest of proteins was performed using Mass Spectrometry Grade Trypsin Gold (Promega, V5280) prepared in 50 mM triethylammonium bicarbonate (TEAB) at a ratio of 1:50 trypsin to protein followed by overnight incubation at 37°C. Resulting peptides were lyophilized by vacuum centrifugation, reconstituted in 2% v/v acetonitrile (ACN)/0.1% v/v formic acid (FA) and adjusted to pH < 3 using 10% FA. Subsequent desalting was performed using Stage Tips prepared using 2× polystyrene-divinylbenzene (SDB-XC) solid-phase extraction disk (Empore, 66884-U) plugs as described with modifications[63]. The SDB-XC membrane was activated using 100% ACN and washed using 2% v/v ACN/0.1% v/v FA. Samples were applied to Stage Tips, centrifuged at 1,800 *g*, and washed using 2% v/v ACN/0.1% v/v FA. Peptides were eluted in 80% v/v ACN/0.1% v/v FA and dried by vacuum centrifugation. Lyophilized peptides were reconstituted in 2% v/v ACN/0.1% v/v FA with the addition of Indexed Retention Time (iRT) peptides (Biognosys, Ki-3002-1) for analysis by LC-MS/MS.

### Mass spectrometry, data processing and analysis

LC-MS/MS was performed on either an Orbitrap Exploris 480 or an Orbitrap Eclipse Tribrid Mass Spectrometer coupled to a Vanquish Neo UHPLC (Thermo Fisher Scientific). Resuspended samples were loaded at a flow rate of 15µL/min onto an Acclaim PepMap 100 trap column (100 µm × 2 cm, nanoViper, C18, 5 µm, 100 Å; Thermo Fisher Scientific). The column was maintained at 40°C. Peptides were eluted from the trap column at a flow rate of 25 µL/min through an Acclaim PepMap RSLC analytical column (75 µm × 50 cm, nanoViper, C18, 2 µm, 100 Å; Thermo Fisher Scientific). HPLC gradient buffer A was 0.1% v/v FA and buffer B was composed of 80% v/v ACN/0.1% FA and was run for 158 min. The mass spectrometer was operated in data-independent acquisition (DIA) mode. For samples run on the Orbitrap Eclipse, MS/MS was run with FAIMS (FAIMS CV: −45 and −60 V). The MS1 Orbitrap resolution for −45 V and −60 V were set to 60 K and 120 K respectively. A full ms1 scan was performed (normalized AGC target: 300%; scan range:380-1000 m/z) and sequential DIA windows (isolation width: 24 m/z) were acquired (scan range: 145 – 1450 m/z; resolution: 30,000; normalized AGC target: 1000%).

DIA analysis was performed using Spectronaut (Version 16.3 or 20.2, Biognosys) using a direct DIA analysis approach. Spectra were searched against the *Homo sapiens* (UP000005640) UniProt FASTA database. Enzyme specificity was set as Trypsin/P and the digest type was specific with 2 missed cleavages allowed. The minimum peptide length was set to 7 and maximum peptide length was 52. Imputing strategy was set as global. Oxidation of methionine and protein N-terminal acetylation were set as variable were set as variable modifications and Carbamidomethylation of cysteines was set as a fixed modification. Raw intensities were exported to Perseus (Version 1.6.15) for filtering of known contaminants, MitoCarta 3.0[3] annotation and statistical analysis using a two-sided Student’s *t*-test. For LFQ, proteins were considered significantly changed where log_2_ (fold change) >1 or <-1 and *p*-value <0.05. For AE-MS, proteins were considered significantly enriched where log_2_ (fold change) >1 and *p*-value <0.05. GO term enrichment analysis was performed using StringDB[64].

### Immunofluorescence

Immunofluorescence was performed as described previously with some modifications[65]. Cells were seeded on #1.5 18×18mm glass coverslips (Carl Zeiss Microscopy) 24-48 hours prior to immunostaining. Specimens were fixed using 4% (w/v) paraformaldehyde in 1× PBS (pH 7.4) for 15 minutes at room temperature and permeabilized using 0.1% v/v Triton X-100 in 1× PBS. Samples were washed three times with 1× PBS followed by blocking with 3% BSA in 1× PBS for 10 minutes. Primary antibodies were prepared in blocking solution, added to coverslips and incubated at room temperature for 2 hours. Primary antibodies and relevant dilutions are listed in **Table S7**. Samples were washed three times with 1× PBS and secondary antibody (Alexa Fluor-488 anti-rabbit IgG, Alexa Fluor-488 anti-mouse IgM, or Alexa Fluor-568 anti-mouse IgG, Thermo Fisher Scientific) was added to coverslips and incubated for 1 hour at room temperature. Where applicable, 10µg/mL of DAPI counterstain was to the secondary antibody staining solution. Following three final washes with 1× PBS, coverslips were mounted onto glass microscope slides (Menzel-Glaser) with mounting medium (90% v/v glycerol, 23µg/mL 1, 4-diazobicyclo-(2,2,2-octane), 100mL Tris-Cl, pH 8.0) and sealed.

### Microscopy

Confocal microscopy was performed using a LSM 980 confocal microscope (Carl Zeiss Microscopy) equipped with 405nm, 488nm, 514nm, 561nm and 639nm lasers, Airyscan 2 detector, 32+2 spectral GaAsP detector with two flanking PMTs and transmitted light PMT TLD using a 63×/1.4 NA oil immersion objective. Z-stacks were acquired at 0.15-0.2µm intervals, 8-10 z-slices per ROI with 2× line averaging and bidirectional line scanning. Images acquired using confocal mode or Airyscan SR mode were post-processed using the Zeiss LSM Plus or Zeiss Airyscan (3D) processing deconvolution algorithm respectively. Representative images were maximum intensity projected and processed using Fiji/ImageJ[66]. For volumetric image analysis, images were acquired using Airyscan 2 Multiplex mode (CO-8Y), 22 0.2µM z-slices per ROI with 0.75× zoom factor, 4× line averaging and bidirectional line scanning. Images acquired in the same experimental group were subject to the same laser power and detector gain settings with some accommodations for variations in fluorescence intensity between samples.

### Electron microscopy

Cell samples for electron microscopy were fixed overnight in Karnovsky’s buffer (2% paraformaldehyde, 2.5% glutaraldehyde in 0.1 M cacodylate buffer, pH 7.4). Subsequent processing was performed in collaboration with the Monash Ramaciotti Centre for Cryo-Electron Microscopy (Cryo-EM), with all steps carried out using the PELCO BioWave system (Ted Pella, Inc.). Samples were post-fixed in 2% (w/v) osmium tetroxide with 1.5% (w/v) potassium ferrocyanide, followed by staining with 1% (w/v) thiocarbohydrazide at 50 °C and a second osmium tetroxide treatment (2% w/v OsO₄ in cacodylate buffer). Milli-Q water washes were performed between each staining step. Cells were then *en bloc* stained overnight in 2% (w/v) aqueous uranyl acetate at 4 °C and subsequently incubated in the BioWave at 50 °C. Finally, samples were stained with freshly prepared Walton’s lead aspartate, washed, and dehydrated through a graded ethanol series (50%, 70%, 90%, 100%), transitioned to acetone, and infiltrated with Hard Epon Procure 812 (Proscitech C045) resin through increasing concentrations in acetone (25%, 50%, 75%, 100%). Beem capsules filled with resin were inverted over the wells and polymerized at 60 °C. Ultrathin sections (70 nm) were cut using a Leica UC7 ultramicrotome (Leica Microsystems) and imaged at 80 kV on a JEOL1400 Plus TEM equipped with a high sensitivity bottom mount CMOS ‘Flash’ camera.

### Image analysis

For mitochondrial volumetric image analysis, organization and data sorting were performed using custom FIJI/ImageJ scripts (v1.54f, NIH, USA). To extract multiple, individual cells from a single volume, the background intensity of the NDUFAF2 staining channel was utilized. The z-stack was initially maximum-intensity projected onto a single plane. Approximately 20 cells from 10 projected planes were manually annotated and used to train a custom Cellpose model using the Cellpose Library from the Medical Imaging Toolbox in MATLAB 2023b[67]. The model was batch applied to the datasets and each respective channel (NDUFAF2, DAPI) was extracted according to the corresponding cell boundary output. Training files were then generated and used to train a segmentation model using the Labkit image pipeline[68]. Both cell masks and mitochondria were then segmented in 3D to generate confidence maps. Confidence maps were passed through MitoLab, a custom in-house developed comprehensive Matlab tool for mitochondrial morphometric analysis. Semantic segmentation was performed by thresholding the confidence maps to generate 3D mitochondrial masks. Binary masks were then processed through a connected-components instance segmentation to separate individual mitochondria with unique identities. Objects smaller than 0.2µm^3^ were filtered out as noise or artefacts. The resulting masks were used to compute morphometric properties. To compute mitochondrial thickness, a distance transformation was performed on an inversely binarized format of the mitochondrial volume. A skeletonized version of the mitochondrial volume was then used to extract the values of the distance transform along the midline of the mitochondrial volume. The final reported value is an average of the entire volume. Mitochondria were classified into four distinct morphological classes: puncta, rod, elongated, and branched using a random forest classifier implemented in Python. A training dataset was curated by manually classifying approximately 700 mitochondria, with roughly 200 of these classified mitochondria specifically selected from the dataset under analysis. Accurate classification for the training set was ensured through a custom-written MATLAB script, which enabled 3D visualization and detailed inspection of mitochondrial morphology. From these 3D images, a set of simple morphometric features was extracted using MATLAB’s Image Processing Toolbox. These basic features were then further processed and combined to create a set of ratio-based morphometric features, including ratios related to surface area to volume, principal axis lengths, and extent. The labelled data was split into training and testing sets using an 80/20 split. The performance of the trained classifier was evaluated by generating confusion matrices. For analysis of nuclear morphology, the Lamin A/C field of views were maximum intensity projected and processed using the Nuclear Irregularity Index plugin in FIJI[69].

### Quantitative PCR (qPCR)

Total mtDNA content was determined by extracting DNA from 120 µg whole-cell protein using a DNeasy Blood and Tissue Kit (QIAGEN, 69506). Primers targeted to the mitochondrial genes *MT-CO1* and *MT-CYB* (*MT-CO1* FWD: 5ʹ-CTCTTCGTCTGATCCGTCCT-3ʹ, *MT-CO1* REV: 5ʹ-TGAGGTTGCGGTCTGTTAGT-3ʹ, *MT-CYB* FWD: 5ʹ-GTAGACAGTCCCACCCTCAC-3ʹ, *MT-CYB* REV: 5ʹ- TTGATCCCGTTTCGTGCAAG-3ʹ) and the nuclear gene *POLG2* (*POLG2* FWD: 5ʹ-CTGCCATAAGGTCTGCAGGT-3ʹ, *POLG2* REV: 5ʹ-CTCCTTTCCGTCAACAGCTC-3ʹ) were used for measurement of mtDNA and nDNA from total DNA extracts. qPCR was performed on a Rotor-Gene Q (QIAGEN, 9001862) using QuantiNova SYBR Green (QIAGEN, 208052) with the following cycling conditions: 95°C for 5 minutes followed by 40 cycles of denaturation (95°C for 15 seconds) and primer annealing and extension (60°C for 40 seconds). Total mtDNA content was determined by normalising the raw Ct value for mtDNA primers to that of nDNA primers (ΔCt). ΔCt values for mtDNA from each genotype were normalized to that of the control (ΔΔCt) and values were expressed as a fold-change.

### Data analysis and statistics

Densitometric analysis of phosphor images was performed using ImageLab 5.2.1 (BioRad) by taking a measure of the signal intensity of radiolabelled species and subtracting the value of the background intensity from each measurement. Signal intensities were normalised to a measure taken of the terminal control time point and expressed as a percentage. Data analysis was performed using GraphPad Prism version 10.6.0 or earlier. All data is expressed as mean +/− SEM. Statistical tests were applied to data as described in figure legends. Comparison plots for discerning unique interactors of each protein were created using RStudio. Custom code was synthesised and modified using Microsoft Copilot and is available upon request.

## Acknowledgements

We thank the members of the Ryan, McArthur and Formosa labs for insightful discussions and research support. We acknowledge funding from the Australian Research Council (DP190103068 to M.T.R.; DP220103559 to M.T.R. and K.M.) and the National Health and Medical Research Council (Ideas Grant #GNT2010939 to M.T.R., EL1 Investigator Grant #GNT2010149 to L.E.F. and Peter Doherty ECF #GNT1161352 to K.M.). S.A. acknowledges funding from the Wellcome Trust Team Science Grant. S.E.J.M. acknowledges the support of the Australian Government Research Training Program scholarship. The authors wish to thank the Monash Proteomics and Metabolomics Facility for provision of instruments and Flowcore (Monash University) for cell-sorting services. We thank Monash Micro Imaging for the provision of equipment, instrument training and insightful discussions. The authors thank the Monash Ramaciotti Centre for Cryo-Electron Microscopy for performing TEM as part of this work and for sharing valuable discussions. The authors thank Micromon Genomics for performing Sanger sequencing of plasmids used in this study. We thank the Biochemistry Imaging Facility (Monash University) for instrument provision, training and technical support. We thank Christina Lackmann for assistance with cloning a MTX1 CRISPR plasmid used in this study. The TOM40 antibody used in this study was a kind gift from Masataka Mori (Kumamoto University).

## Author contributions

(Using CRediT Taxonomy)

<u>Sarah E. J. Morf:</u> Conceptualization, data curation, formal analysis, investigation, methodology, project administration, validation, visualization, writing – original draft preparation

<u>Matthew P. Challis:</u> Data curation, formal analysis, methodology, investigation, visualization

<u>Sanjeev Uthishtran:</u> Data curation, formal analysis, methodology, resources, software

<u>Caitlin L. Rowe:</u> Investigation, methodology

<u>Alice J. Sharpe:</u> Investigation, methodology

<u>Natasha Kapoor-Kaushik:</u> Investigation, methodology

<u>Senthil Arumugam:</u> Methodology, resources, supervision

<u>Luke E. Formosa:</u> Conceptualization, methodology, project administration, resources, supervision, writing – original draft preparation

<u>Kate McArthur:</u> Methodology, funding acquisition, project administration, resources, supervision, writing – original draft preparation

<u>Michael T. Ryan:</u> Conceptualization, funding acquisition, project administration, resources, supervision, visualization, writing – original draft preparation

All authors contributed to writing – review and editing.

## References

1. McBride HM, Neuspiel M, Wasiak S. Mitochondria: More Than Just a Powerhouse. Curr Biol. 2006. DOI: 10.1016/j.cub.2006.06.054

2. Spinelli JB, Haigis MC. The multifaceted contributions of mitochondria to cellular metabolism. Nat Cell Biol. 2018. DOI: 10.1038/s41556-018-0124-1

3. Rath S, Sharma R, Gupta R, Ast T, Chan C, Durham TJ, et al. MitoCarta3.0: an updated mitochondrial proteome now with sub-organelle localization and pathway annotations. Nucleic Acids Res. 2021. DOI: 10.1093/nar/gkaa1011

4. Chacinska A, Koehler CM, Milenkovic D, Lithgow T, Pfanner N. Importing Mitochondrial Proteins: Machineries and Mechanisms. Cell. 2009. DOI: 10.1016/j.cell.2009.08.005

5. Pfanner N, Warscheid B, Wiedemann N. Mitochondrial proteins: from biogenesis to functional networks. Nat Rev Mol Cell Biol. 2019. DOI: 10.1038/s41580-018-0092-0

6. Hill K, Model K, Ryan MT, Dietmeier K, Martin F, Wagner R, et al. Tom40 forms the hydrophilic channel of the mitochondrial import pore for preproteins. Nature. 1998. DOI: 10.1038/26780

7. Höhr AIC, Straub SP, Warscheid B, Becker T, Wiedemann N. Assembly of β-barrel proteins in the mitochondrial outer membrane. BBA Mol Cell Res. 2015. DOI: 10.1016/j.bbamcr.2014.10.006

8. Kozjak V, Wiedemann N, Milenkovic D, Lohaus C, Meyer HE, Guiard B, et al. An essential role of Sam50 in the protein sorting and assembly machinery of the mitochondrial outer membrane. J Biol Chem. 2003. DOI: 10.1074/jbc.C300442200

9. Diederichs KA, Buchanan SK, Botos I. Building Better Barrels – β-barrel Biogenesis and Insertion in Bacteria and Mitochondria. J Mol Biol. 2021. 10.1016/j.jmb.2021.166894

10. Morgenstern M, Peikert CD, Lübbert P, Suppanz I, Klemm C, Alka O, et al. Quantitative high-confidence human mitochondrial proteome and its dynamics in cellular context. Cell Metab. 2021. DOI: 10.1016/j.cmet.2021.11.001

11. Diederichs KA, Ni X, Rollauer SE, Botos I, Tan X, King MS, et al. Structural insight into mitochondrial β-barrel outer membrane protein biogenesis. Nat Commun. 2020. DOI: 10.1038/s41467-020-17144-1

12. Takeda H, Tsutsumi A, Nishizawa T, Lindau C, Busto J V, Wenz LS, et al. Mitochondrial sorting and assembly machinery operates by β-barrel switching. Nature. 2021. DOI: 10.1038/s41586-020-03113-7

13. Huynen MA, Mühlmeister M, Gotthardt K, Guerrero-Castillo S, Brandt U. Evolution and structural organization of the mitochondrial contact site (MICOS) complex and the mitochondrial intermembrane space bridging (MIB) complex. BBA Mol Cell Res. 2016. DOI: 10.1016/j.bbamcr.2015.10.009

14. Elouej S, Harhouri K, Mao M Le, Baujat G, Nampoothiri S, Kayserili HU, et al. Loss of MTX2 causes mandibuloacral dysplasia and links mitochondrial dysfunction to altered nuclear morphology. Nat Commun. 2020. DOI: 10.1038/s41467-020-18146-9

15. Doğan BY, Günay N, Ada Y, Doğan ME. A novel MTX2 gene splice site variant resulting in exon skipping, causing the recently described mandibuloacral dysplasia progeroid syndrome. Am J Med Genet A. 2023. DOI: 10.1002/ajmg.a.63010

16. Ding C, Wu Z, Huang L, Wang Y, Xue J, Chen S, et al. Mitofilin and CHCHD6 physically interact with Sam50 to sustain cristae structure. Sci Rep. 2015. DOI: 10.1038/srep16064

17. Ott C, Ross K, Straub S, Thiede B, Gotz M, Goosmann C, et al. Sam50 Functions in Mitochondrial Intermembrane Space Bridging and Biogenesis of Respiratory Complexes. Mol Cell Biol. 2012. DOI: 10.1128/mcb.06388-11

18. Qu Y, Jelicic B, Pettolino F, Perry A, Lo TL, Hewitt VL, et al. Mitochondrial sorting and assembly machinery subunit Sam37 in Candida albicans: Insight into the roles of mitochondria in fitness, cell wall integrity, and virulence. Eukaryot Cell. 2012. DOI: 10.1128/EC.05292-11

19. Kozjak-Pavlovic V, Ross K, Benlasfer N, Kimmig S, Karlas A, Rudel T. Conserved roles of Sam50 and metaxins in VDAC biogenesis. EMBO Rep. 2007. DOI: 10.1038/sj.embor.7400982

20. Humphries AD, Streimann IC, Stojanovski D, Johnston AJ, Yano M, Hoogenraad NJ, et al. Dissection of the mitochondrial import and assembly pathway for human Tom40. J Biol Chem. 2005. DOI: 10.1074/jbc.M413816200

21. Ashburner M, Ball CA, Blake JA, Botstein D, Butler H, Cherry JM, et al. Gene ontology: Tool for the unification of biology. Nat Genet. 2000. DOI: 10.1038/75556

22. Shembekar SS, Nikolaus P, Honnert U, Höring M, Attia A, Topp K, et al. Regulation of mitochondrial cristae organization by Myo19, Miro1/2 and Metaxin 3. J Cell Sci. 2025. DOI: 10.1242/jcs.263637

23. Ioakeimidis F, Ott C, Kozjak-Pavlovic V, Violitzi F, Rinotas V, Makrinou E, et al. A splicing mutation in the novel mitochondrial protein DNAJC11 causes motor neuron pathology associated with cristae disorganization, and lymphoid abnormalities in mice. PLoS One. 2014. DOI: 10.1371/journal.pone.0104237

24. Abdul KM, Terada K, Yano M, Ryan MT, Streimann I, Hoogenraad NJ, et al. Functional analysis of human metaxin in mitochondrial protein import in cultured cells and its relationship with the Tom complex. Biochem Biophys Res Commun. 2000. DOI: 10.1006/bbrc.2000.3589

25. Wagner F, Kunz TC, Chowdhury SR, Thiede B, Fraunholz M, Eger D, et al. Armadillo repeat-containing protein 1 is a dual localization protein associated with mitochondrial intermembrane space bridging complex. PLoS One. 2019. DOI: 10.1371/journal.pone.0218303

26. Gok MO, Connor OM, Wang X, Menezes CJ, Llamas CB, Mishra P, et al. The outer mitochondrial membrane protein TMEM11 demarcates spatially restricted BNIP3/BNIP3L-mediated mitophagy. J of Cell Biol. 2023. DOI: 10.1083/jcb.202204021

27. McKenna MJ, Kraus F, Coelho JPL, Vasandani M, Zhang J, Adams BM, et al. ARMC1 partitions between distinct complexes and assembles MIRO with MTFR to control mitochondrial distribution. Sci Adv. 2025. DOI: 10.1126/sciadv.adu5091

28. Rios KE, Zhou M, Lott NM, Beauregard CR, McDaniel DP, Conrads TP, et al. CARD19 Interacts with Mitochondrial Contact Site and Cristae Organizing System Constituent Proteins and Regulates Cristae Morphology. Cells. 2022. DOI: 10.3390/cells11071175

29. Gandre-Babbe S, van der Bliek AM. The Novel Tail-anchored Membrane Protein Mff Controls Mitochondrial and Peroxisomal Fission in Mammalian Cells. Mol Biol Cell. 2008. DOI: 10.1091/mbc.e07-12-1287

30. Osellame LD, Singh AP, Stroud DA, Palmer CS, Stojanovski D, Ramachandran R, et al. Cooperative and independent roles of the Drp1 adaptors Mff, MiD49 and MiD51 in mitochondrial fission. J Cell Sci. 2016. DOI: 10.1242/jcs.185165

31. Tilokani L, Russell FM, Hamilton S, Virga DM, Segawa M, Paupe V, et al. AMPK-dependent phosphorylation of MTFR1L regulates mitochondrial morphology. Sci Adv. 2022. DOI: 10.1126/sciadv.abo7956

32. Zhang T, Li L, Fan X, Shou X, Ruan Y, Xie X. Metaxin-2 tunes mitochondrial transportation and neuronal function in *Drosophila*. Genetics. 2024. DOI: 10.1093/genetics/iyae204

33. Zhao Y, Song E, Wang W, Hsieh CH, Wang X, Feng W, et al. Metaxins are core components of mitochondrial transport adaptor complexes. Nat Commun. 2021. DOI: 10.1038/s41467-020-20346-2

34. Shi P, Ren X, Meng J, Kang C, Wu Y, Rong Y, et al. Mechanical instability generated by Myosin 19 contributes to mitochondria cristae architecture and OXPHOS. Nat Commun. 2022. DOI: 10.1038/s41467-022-30431-3

35. Oeding SJ, Majstrowicz K, Hu XP, Schwarz V, Freitag A, Honnert U, et al. Identification of Miro1 and Miro2 as mitochondrial receptors for myosin XIX. J Cell Sci. 2018. DOI: 10.1242/jcs.219469

36. López-Doménech G, Covill-Cooke C, Ivankovic D, Halff EF, Sheehan DF, Norkett R, et al. Miro proteins coordinate microtubule- and actin-dependent mitochondrial transport and distribution. EMBO J. 2018. DOI: 10.15252/embj.201696380

37. Szklarczyk R, Huynen MA. Expansion of the human mitochondrial proteome by intra- and inter-compartmental protein duplication. Genome Biol. 2009. DOI: 10.1186/gb-2009-10-11-r135

38. Roger AJ, Muñoz-Gómez SA, Kamikawa R. The Origin and Diversification of Mitochondria. Curr Biol. 2017. DOI: 10.1016/j.cub.2017.09.015

39. Weinhäupl K, Lindau C, Hessel A, Wang Y, Schütze C, Jores T, et al. Structural Basis of Membrane Protein Chaperoning through the Mitochondrial Intermembrane Space. Cell. 2018. DOI: 10.1016/j.cell.2018.10.039

40. Kutik S, Stojanovski D, Becker L, Becker T, Meinecke M, Krüger V, et al. Dissecting Membrane Insertion of Mitochondrial β-Barrel Proteins. Cell. 2008. DOI: 10.1016/j.cell.2008.01.028

41. Wang Q, Guan Z, Qi L, Zhuang J, Wang C, Hong S, et al. Structural insight into the SAM-mediated assembly of the mitochondrial TOM core complex. Science. 2021. DOI: 10.1126/science.abh0704

42. Armstrong LC, Komiya T, Bergman BE, Mihara K, Bornstein P. Metaxin is a component of a preprotein import complex in the outer membrane of the mammalian mitochondrion. J Biol Chem. 1997. DOI: 10.1074/jbc.272.10.6510

43. Armstrong LC, Saenz AJ, Bornstein P. Metaxin 1 interacts with metaxin 2, a novel related protein associated with the mammalian mitochondrial outer membrane. J Cell Biochem. 1999. DOI: 10.1002/(SICI)1097-4644(19990701)74:1<11::AID-JCB2>3.0.CO;2-V

44. Xie J, Marusich MF, Souda P, Whitelegge J, Capaldi RA. The mitochondrial inner membrane protein Mitofilin exists as a complex with SAM50, metaxins 1 and 2, coiled-coil-helix coiled-coil-helix domain-containing protein 3 and 6 and DnaJC11. FEBS Lett. 2007. DOI: 10.1016/j.febslet.2007.06.052

45. Guarani V, McNeill EM, Paulo JA, Huttlin EL, Fröhlich F, Gygi SP, et al. QIL1 is a novel mitochondrial protein required for MICOS complex stability and cristae morphology. Elife. 2015. DOI: 10.7554/eLife.06265

46. Antonicka H, Lin ZY, Janer A, Aaltonen MJ, Weraarpachai W, Gingras AC, et al. A High-Density Human Mitochondrial Proximity Interaction Network. Cell Metab. 2020. DOI: 10.1016/j.cmet.2020.07.017

47. Coscia SM, Thompson CP, Tang Q, Baltrusaitis EE, Rhodenhiser JA, Quintero-Carmona OA, et al. Myo19 tethers mitochondria to endoplasmic reticulum-associated actin to promote mitochondrial fission. J Cell Sci. 2023. DOI: 10.1242/jcs.260612

48. Morciano G, Naumova N, Koprowski P, Valente S, Sardão VA, Potes Y, et al. The mitochondrial permeability transition pore: an evolving concept critical for cell life and death. Biol Rev. 2021. DOI: 10.1111/brv.12764

49. Bernardi P, Gerle C, Halestrap AP, Jonas EA, Karch J, Mnatsakanyan N, et al. Identity, structure, and function of the mitochondrial permeability transition pore: controversies, consensus, recent advances, and future directions. Cell Death Differ. 2023. DOI: 10.1038/s41418-023-01187-0

50. Yamada T, Ikeda A, Murata D, Wang H, Zhang C, Khare P, et al. Dual regulation of mitochondrial fusion by Parkin–PINK1 and OMA1. Nature. 2025. DOI: 10.1038/s41586-025-08590-2

51. Yamada T, Murata D, Adachi Y, Itoh K, Kameoka S, Igarashi A, et al. Mitochondrial Stasis Reveals p62-Mediated Ubiquitination in Parkin-Independent Mitophagy and Mitigates Nonalcoholic Fatty Liver Disease. Cell Metab. 2018. DOI: 10.1016/j.cmet.2018.06.014

52. Ma X, Chen A, Melo L, Clemente-Sanchez A, Chao X, Ahmadi AR, et al. Loss of hepatic DRP1 exacerbates alcoholic hepatitis by inducing megamitochondria and mitochondrial maladaptation. Hepatology. 2023. DOI: 10.1002/hep.32604

53. Ban-Ishihara R, Ishihara T, Sasaki N, Mihara K, Ishihara N. Dynamics of nucleoid structure regulated by mitochondrial fission contributes to cristae reformation and release of cytochrome c. Proc Natl Acad Sci U S A. 2013. DOI: 10.1073/pnas.1301951110

54. Kleele T, Rey T, Winter J, Zaganelli S, Mahecic D, Lambert HP, et al. Distinct fission signatures predict mitochondrial degradation or biogenesis. Nature. 2021. DOI: 10.1038/s41586-021-03510-6

55. Garg A, Keng WT, Chen Z, Sathe AA, Xing C, Kailasam PD, et al. Autosomal recessive progeroid syndrome due to homozygosity for a TOMM7 variant. J Clin Invest. 2022. DOI: 10.1172/JCI156864

56. Young C, Batkovskyte D, Kitamura M, Shvedova M, Mihara Y, Akiba J, et al. A hypomorphic variant in the translocase of the outer mitochondrial membrane complex subunit TOMM7 causes short stature and developmental delay. HGG Adv. 2023. DOI: 10.1016/j.xhgg.2022.100148

57. Labun K, Montague TG, Krause M, Cleuren YNT, Tjeldnes H, Valen E. CHOPCHOP v3: Expanding the CRISPR web toolbox beyond genome editing. Nucleic Acids Res. 2019. DOI: 10.1093/nar/gkz365

58. Ran FA, Hsu PD, Wright J, Agarwala V, Scott DA, Zhang F. Genome engineering using the CRISPR-Cas9 system. Nat Protoc. 2013. DOI: 10.1038/nprot.2013.143

59. Formosa LE, Muellner-Wong L, Reljic B, Sharpe AJ, Jackson TD, Beilharz TH, et al. Dissecting the Roles of Mitochondrial Complex I Intermediate Assembly Complex Factors in the Biogenesis of Complex I. Cell Rep. 2020. DOI: 10.1016/j.celrep.2020.107541

60. Johnston AJ, Hoogenraad J, Dougan DA, Truscott KN, Yano M, Mori M, et al. Insertion and assembly of human Tom7 into the preprotein translocase complex of the outer mitochondrial membrane. J Biol Chem. 2002. DOI: 10.1074/jbc.M205613200

61. Schägger H, von Jagow G. Blue native electrophoresis for isolation of membrane protein complexes in enzymatically active form. Anal Biochem. 1991. DOI: 10.1016/0003-2697(91)90094-A

62. Schägger H, von Jagow G. Tricine-sodium dodecyl sulfate-polyacrylamide gel electrophoresis for the separation of proteins in the range from 1 to 100 kDa. Anal Biochem. 1987. DOI: 10.1016/0003-2697(87)90587-2

63. Rappsilber J, Ishihama Y, Mann M. Stop and Go Extraction Tips for Matrix-Assisted Laser Desorption/Ionization, Nanoelectrospray, and LC/MS Sample Pretreatment in Proteomics. Anal Chem. 2003. DOI: 10.1021/ac026117i

64. Szklarczyk D, Kirsch R, Koutrouli M, Nastou K, Mehryary F, Hachilif R, et al. The STRING database in 2023: protein-protein association networks and functional enrichment analyses for any sequenced genome of interest. Nucleic Acids Res. 2023. DOI: 10.1093/nar/gkac1000

65. Kamerkar SC, Kraus F, Sharpe AJ, Pucadyil TJ, Ryan MT. Dynamin-related protein 1 has membrane constricting and severing abilities sufficient for mitochondrial and peroxisomal fission. Nat Commun. 2018. DOI: 10.1038/s41467-018-07543-w

66. Schindelin J, Arganda-Carreras I, Frise E, Kaynig V, Longair M, Pietzsch T, et al. Fiji: An open-source platform for biological-image analysis. Nat Methods. 2012. DOI: 10.1038/nmeth.2019

67. Pachitariu M, Stringer C. Cellpose 2.0: how to train your own model. Nat Methods. 2022. DOI: 10.1038/s41592-022-01663-4

68. Arzt M, Deschamps J, Schmied C, Pietzsch T, Schmidt D, Tomancak P, et al. LABKIT: Labeling and Segmentation Toolkit for Big Image Data. Front Comput Sci. 2022. DOI: 10.3389/fcomp.2022.777728

69. Svoren M, Camerini E, van Erp M, Yang FW, Bakker G-J, Wolf K. Approaches to Determine Nuclear Shape in Cells During Migration Through Collagen Matrices. 2023. p. 97–114. DOI: 10.1007/978-1-0716-2887-4_7

70. Formosa LE, Mimaki M, Frazier AE, McKenzie M, Stait TL, Thorburn DR, et al. Characterization of mitochondrial FOXRED1 in the assembly of respiratory chain complex I. Hum Mol Genet. 2015. DOI: 10.1093/hmg/ddv058

