## Supplementary File for "Characterization of human Metaxin proteins reveals functional diversification of SAM37 homologs MTX1 and MTX3"

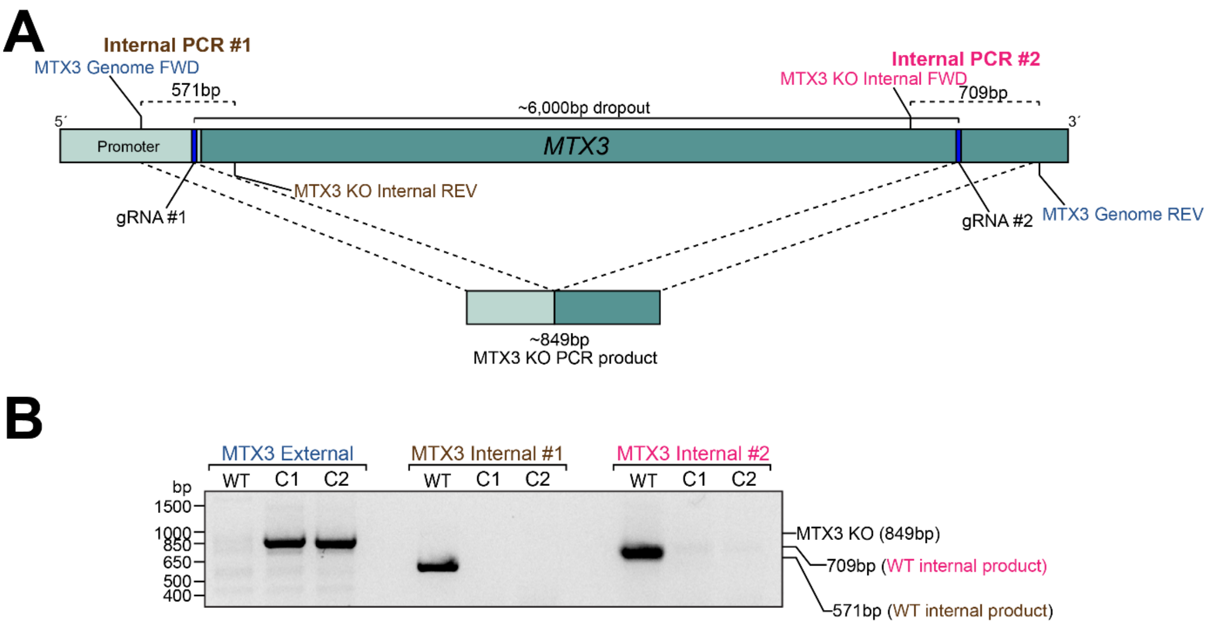


**Fig. S1. Validation of MTX3$^{KO}$ cells by genotyping. A).** Schematic illustrating CRISPR target sites and primer design for validating MTX3$^{KO}$ cells by genotyping (not to scale). Dark blue bands represent approximate location of CRISPR targeting sites. Blue text denotes primer binding sites for "external" genotyping primers. Brown and magenta text indicate position of "internal" primer binding sites. **B).** PCR product amplified from gDNA extracted from MTX3$^{KO}$ clone 1 (C1) and clone 2 (C2) using selected primer combinations. Presence of an 849bp product in MTX3$^{KO}$ C1 and C2 indicates successful excision of ~6,000bp sequence between CRISPR target sites. Absence of PCR product amplified using internal primer sets in MTX3$^{KO}$ C1 and C2 indicates removal of primer binding site by successful excision of the *MTX3* gene sequence.

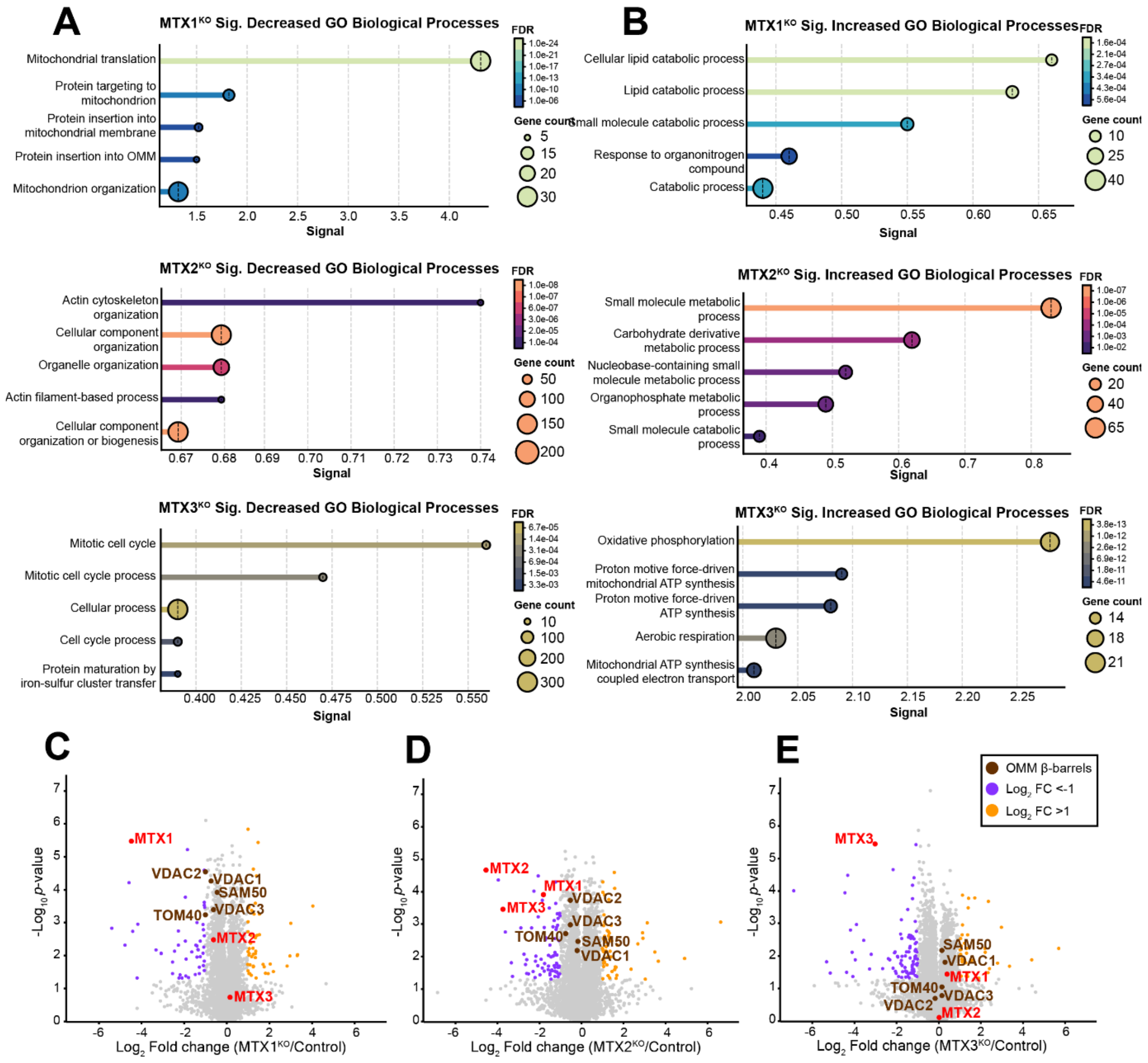


**Figure S2. Changes to the whole-cell proteomic landscape in cells lacking MTX1, MTX2 or MTX3.** Top 5 Gene Ontology (GO) Biological Processes ranked by signal strength of **A).** significantly decreased proteins with a $Log_2$ FC <-0.5 and *p*-value <0.05 or **B).** significantly increased proteins with a $Log_2$ FC >0.5 and *p*-value <0.05 (n=3 biological replicates, two-sided Students' *t*-test) Functional enrichment analysis was performed using string-db.org. Volcano plots highlighting changes to known OMM β-barrel proteins in the absence of **C).** MTX1, **D).** MTX2, or **E).** MTX3.

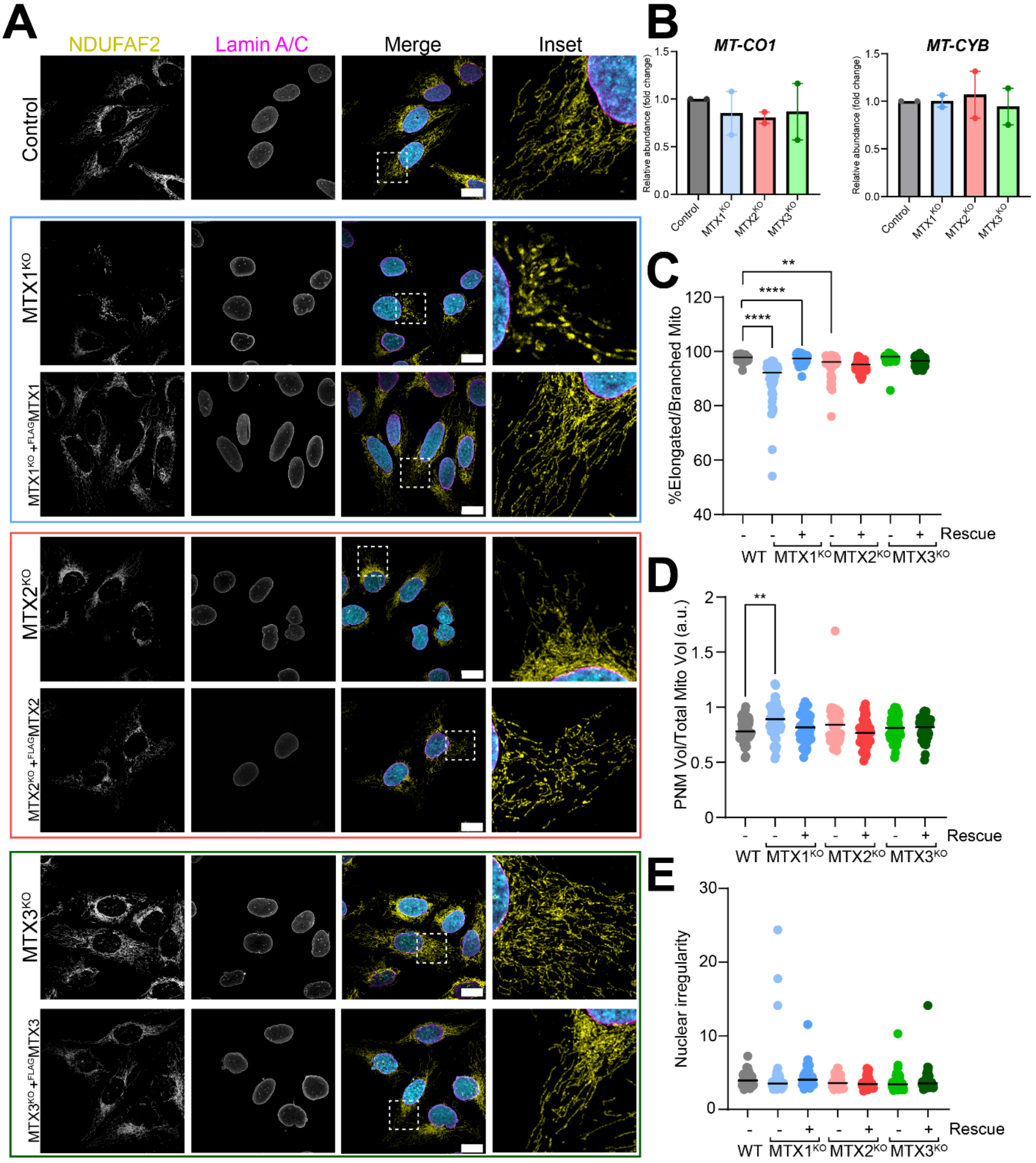


**Figure S3. Changes to mitochondrial morphology and mtDNA distribution resulting from loss of Metaxins. A).** Representative confocal images of control, $MTX1^{KO}$, $MTX1^{KO}$ + $^{FLAG}MTX1$, $MTX2^{KO}$, $MTX2^{KO}$ + $^{FLAG}MTX2$, $MTX3^{KO}$ and $MTX3^{KO}$ + $^{FLAG}MTX3$ cells at steady-state. Immunofluorescence was performed using antibodies against NDUFAF2 (matrix, yellow), and LaminA/C (nuclear envelope, magenta) with a DAPI counterstain (nuclear DNA, cyan). Scale bar = 20µm. **B).** Quantification of total mtDNA content measured by qPCR. Volumetric quantification of **C).** % of mitochondria classified as “branched” or “elongated”, **D).** volume of mitochondria occupying the perinuclear area of the cell as a proportion of total mitochondria according to NDUFAF2 staining and **E).** nuclei quantified for nuclear irregularity according to Lamin A/C staining in control (n=50 cells), $MTX1^{KO}$ (n=56 cells), $MTX1^{KO}$ + $^{FLAG}$MTX1 (n=52 cells), $MTX2^{KO}$ (n=47 cells), $MTX2^{KO}$ + $^{FLAG}$MTX2 (n=50 cells), $MTX3^{KO}$ (n=47 cells) and $MTX3^{KO}$ + $^{FLAG}$MTX3 (n=47 cells) cells. All statistical tests were performed using an ordinary one-way ANOVA with Tukey's multiple comparisons test (ns $p>0.05$, * $p<0.05$, ** $p<0.01$, **** $p<0.0001$).

**Supplementary table 5.** Targeted gRNA sequences for CRISPR/Cas9-mediated genome editing

| **Uniprot ID** | **Gene** | | **gRNA sequence PAM (5′-3′)** | **Target** |
|---|---|---|---|---|
| Q13505-3 | *MTX1* (Isoform 3) | **1** | GCCGCGCCTTCAGGGGTTCG | Intronic region upstream of exon 1 (sense strand) |
| | | **2** | AAGAGCGTACGTGGCCGATATGG | Intronic region downstream of exon 1 (sense strand) |
| O75431-1 | *MTX2* (Isoform 1) | **1** | CGGCACTCGAGCGCTTCCGCCGG | Intronic region upstream of exon 1 (antisense strand) |
| | | **2** | GTTCAGCGTATAATGCCGCAAGG | Intronic region downstream of exon 1 (antisense strand) |
| Q5HYI7-1 Q5HYI7-4 Q5HYI7-5 | *MTX3* (multiple isoforms) | **1** | TTTCCGGCACGTGAAACGCGGGG | Intronic region upstream of exon 1 (antisense strand) |
| | | **2** | CAGTTAGTACCCTTTAAGCTAGG | Intronic region downstream of exon 9, isoform 1 (sense strand) |

**Supplementary table 6.** Genotyping primers for validation of targeted gene disruption

| **Target gene** | **Primer name** | **Sequence (5′-3′)** |
|---|---|---|
| *MTX1* | MTX1 Genome FWD | GAAAGGGACACCAGGGACAG |
| | MTX1 Genome REV | GGCTGTGAGTCGTTTGGTCT |
| | MTX1 Internal REV | GACCAGCAGAACAGCTCCAT |
| *MTX2* | MTX2 Genome FWD | TGGCTTACAGAAAGGACGCT |
| | MTX2 Genome REV | GGAGGTGGGACAAAGTACCG |
| | MTX2 Internal REV | TGCAATCTGGGAGACGAAGG |
| *MTX3* | MTX3 Genome FWD | AAGGACCAGCACTGAAGGTG |
| | MTX3 Genome REV | AGAGAGGCAGACACACAAGC |
| | MTX3 Internal FWD | CCAGCTGGACAAGAAACGGT |
| | MTX3 Internal REV | TTTCGAGGACTTTGCTCCCC |

**Supplementary table 7.** Antibodies used in this study.

| Target | Host | Working Dilution | Diluent | Source (Product no.) |
|---|---|---|---|---|
| MTX1 | Mouse | 1:500 (WB) | 3% BSA, 1× TBS, 0.1% Tween-20 | Santa Cruz Biotechnology (sc-514469) |
| MTX2 | Mouse | 1:1000 (WB) | 3% BSA, 1× TBS, 0.1% Tween-20 | Santa Cruz Biotechnology (sc-514231) |
| FLAG | Mouse | 1:1000 (WB) | 3% BSA, 1× TBS, 0.1% Tween-20 | Sigma-Aldrich (F1804) |
| SAM50 | Rabbit | 1:500 (WB) | 5% skim milk, 1× PBS | Ryan Lab In-House[20] (N/A) |
| SDHA | Mouse | 1:1000 (WB) | 3% BSA, 1× TBS, 0.1% Tween-20 | Abcam (ab14715) |
| VDAC1 | Rabbit | 1:500 (WB) | 5% skim milk, 1× TBS, 0.1% Tween-20 | Ryan Lab In-House (N/A) |
| TOM40 | Rabbit | 1:500 (WB) | 5% skim milk, 1× TBS, 0.1% Tween-20 | Gift from M. Mori (Kumamoto University)[24] |
| HSP60 | Mouse | 1:500 (IFA) | 3% BSA, 1× PBS | Abcam (ab46798) |
| DNA | Mouse | 1:200 (IFA) | 3% BSA, 1× PBS | Progen (AC-30-10) |
| NDUFAF2 | Rabbit | 1:500 (IFA) | 3% BSA, 1× PBS | Ryan Lab In-House[70] (N/A) |
| Lamin A/C | Mouse | 1:500 (IFA) | 3% BSA, 1× PBS | Santa Cruz Biotechnology (sc-376248) |
